# Guava *cv.* Allahabad Safeda Chromosome scale assembly and comparative genomics decodes breeders’ choice marker trait association for pink pulp colour

**DOI:** 10.1101/2024.03.29.587319

**Authors:** Amandeep Mittal, Sujata Thakur, Ankush Sharma, Rajbir Singh Boora, Naresh Kumar Arora, Daljinder Singh, Manav Indra Singh Gill, Guriqbal Singh Dhillon, Parveen Chhuneja, Inderjit Singh Yadav, Manish Jindal, Oommen K. Mathew, Vijaya Bhasker Reddy Lachagari, Andrew H. Paterson

**Affiliations:** School of Agricultural Biotechnology, Punjab Agricultural University, Ludhiana, Punjab 141004, India; Fruit Research Sub-Station, Punjab Agricultural University, Bahadurgarh, Patiala 147002, India; Department of Fruit Science, Punjab Agricultural University, Ludhiana 141004, India; AgriGenome Labs Private Ltd, Kochi 682042, Kerala, India – Current Affiliation - MedGenome Labs, Hosur Rd, Bengaluru, Karnataka 560099; AgriGenome Labs Private Ltd, Genome Valley, Shamirpet 500078, Hyderabad, India - Current Affiliation - ATGC Biotech Pvt Ltd, ATGC innovation Square, Genome Valley, Biotech Park, Hyderabad 500 078; Plant Genome Mapping Laboratory, University of Georgia, Athens, Georgia 30605, USA

**Keywords:** *Psidium guajava* L., Genome assembly, Genome re-sequencing, Genotyping by sequencing - ddRAD, Association mapping, Pulp colour, Marker assisted breeding

## Abstract

Deciphering chromosomal length genome assemblies has the potential to unravel an organism’s evolutionary relationships and genetic mapping of traits of commercial importance. We assembled guava genome using a hybrid sequencing approach with ∼450x depth Illumina short reads, ∼35x PacBio long reads and Bionano maps to ∼594 MB Scaffold length on 11 pseudo chromosomes (∼479 MB contig length). Maker pipeline predicted 17,395 genes, 23% greater from earlier draft produced in same cultivar Allahabad Safeda. The genome assembly clarified guava evolutionary history, for example revealing predominance of gene expansion by dispersed duplications, in particular contributing to abundance of monoterpene synthases; and supporting evidence of a whole genome duplication event in guava as in other Myrtaceae. Guava breeders have been aiming to reduce screening time for selecting pink pulp colour progenies using marker-trait associations, but a previous comparative transcriptomics and comparative genomics approach with draft genome assembly to identify the effector gene associated with pink pulp was unsuccessful. Here, genome re-sequencing with Illumina short reads at ∼25x depth of 20 pink fleshed and/or non-coloured guava cultivars and comprehensive analysis for genes in the carotenoid biosynthesis pathway identified structural variations in *Phytoene Synthase* 2. Further, ddRAD based association mapping in core-collection of 82 coloured and non-coloured genotypes from Indian sub-continent found strong association with the same causal gene. Subsequently, we developed PCR based Indel/SSR breeder friendly marker that can readily be scored in routine agarose gels and empowers accurate selection for seedlings that will produce fruits with pink pulp.

## Introduction

Guava (*Psidium guajava* L.), a perennial tree native to central America, is an often-cross pollinated plant belonging to the family Myrtaceae with chromosome number 2n=22. The genome size of guava is approximately 465 Mbp (Feuillet et al., 2011) although larger genome size of 495-538 MB has been reported for Brazilian white and red-fleshed diploid cultivars. Tetraploid *P. cattleianum* and *P. acutangulum* have been reported to have still larger genomes of 1030 -1144 MB (Da Costa et al., 2008). Guava is known as ‘apple of the tropics’ due to its low cultivation cost, and is grown in tropical and sub-tropical regions of the world. It is an economically significant crop in countries such as India, China, Pakistan, Mexico, Brazil, Egypt, Indonesia, Columbia, and Thailand (Gangappa et al., 2022). The fruit is a rich source of nutrients (Thaipong et al., 2006; Mittal et al., 2020) including minerals, vitamins, phosphorus, iron, calcium, zinc, and vitamin C. One peculiar feature of guava fruit is its variability in fruit pulp color with pink and white pulp being the most prominent types(Mehmood et al., 2014).

Traditional targeted breeding in fruit crops is time-consuming, expensive, and labour-intensive owing to the years-long juvenile phase(Longhi et al., 2013) delaying the process of fruit evaluation and characterization. In the era of next generation sequencing (NGS) these constraints can be overcome by chromosome landing by association mapping, but decoding of genomes is required for strong associations. Shotgun sequencing approaches and strong computational algorithms mitigate the need for massively expensive and consortia-based clone to contig approaches. Over the past decade, more than 50 fruit and vegetable genomes have been sequenced and assembled to chromosomal scale using shotgun sequencing. Massively parallel re-sequencing of smaller genomes and screening for candidate genes in contrasting trait genotypes has the great potential for identification of trait specific markers. Rapid genotyping by sequencing (GBS) approaches like double digest - Restriction site Associated DNA Sequencing (ddRAD-Seq) can cheaply identify and validate new marker-trait associations. The associated genomic islands developed into simple PCR-based diagnostic markers enable affordable marker-assisted breeding (MAB)(Baumgartner et al., 2016)(Varshney et al., 2005; Kole et al., 2015) in segregating populations, reducing much in-field testing to in-lab assays.

Carotenoids are tetraterpene natural pigments providing red, orange, or yellow colour to leaves, flowers, fruits and vegetables (Gupta and Hirschberg, 2022a)(Ma et al., 2023), also conferring antioxidant properties in plants(Jang et al., 2020). During biosynthesis, two Geranylgeranyl diphosphate (GGPP) molecules condense to form phytoene in the presence of a rate-limiting regulatory enzyme, phytoene synthase (PSY). This conversion is the first step of the pathway that subsequently leads to the formation of lycopene which further gets converted to carotenes (Gupta and Hirschberg, 2022b). PSY’s role in carotenoid accumulation has been reported in several loss or gain of function studies where changes in the functional copy of PSY led to altered accumulation of carotenoid pigments. PSY mutants of carrots display pale pigmentation in callus (Oleszkiewicz et al., 2021). Similar impacts have been reported in tomato, maize and wheat where mutation in PSY led to reduced accumulation of carotenoids (Zhu et al., 2016; Chen et al., 2019; Zhang et al., 2021). There is a single PSY gene in Arabidopsis(Lindgren et al., 2003) whereas two or more isoforms have been reported in tomato(Chen et al., 2019), citrus(Ma et al., 2023), carrot(Just et al., 2007; Bowman et al., 2014), capsicum(Jang et al., 2020), melon(Qin et al., 2011), loquat(Fu et al., 2012), rice(Welsch et al., 2008).

Here, we report genome scale assembly of commercially important Indian guava cultivar Allahabad Safeda (AS). AS has been widely cultivated commercially for over 5 decades in India owing to its desirable fruit characteristics including attractive fruit size, excellent organoleptic traits, and yield stability across diverse agroclimatic zones. Despite the widespread use of AS in breeding programs, the Indian guava germplasm remains largely unexplored, and no trait-specific diagnostic markers have been discovered to date. Previously, we attempted to identify the candidate genes for pink pulp colour in guava by comparative transcriptomics of white and pink pulp cultivars(Mittal et al., 2020) but the complexity of lycopene and ethylene signalling prevented the identification of any strong association. However, by utilizing a chromosomal scale genome assembly, re-sequencing of 20 core pulp colour variable genotypes and genome wide association mapping in a panel of 82 genotypes, we identified a marker scorable on agarose gels that differentiates pink from white pulp in the Indian nation-wide guava germplasm. This discovery could aid in the development of affordable and efficient marker-assisted breeding strategies for guava, facilitating the improvement of this important crop.

## Results

### Genome Assembly

The High Molecular Weight (HMW) DNA with 150-200 KB average fragment length was extracted from etiolated leaves of a ∼10-year-old guava cv. Allahabad Safeda tree (Figure1A).

**Figure 1.**
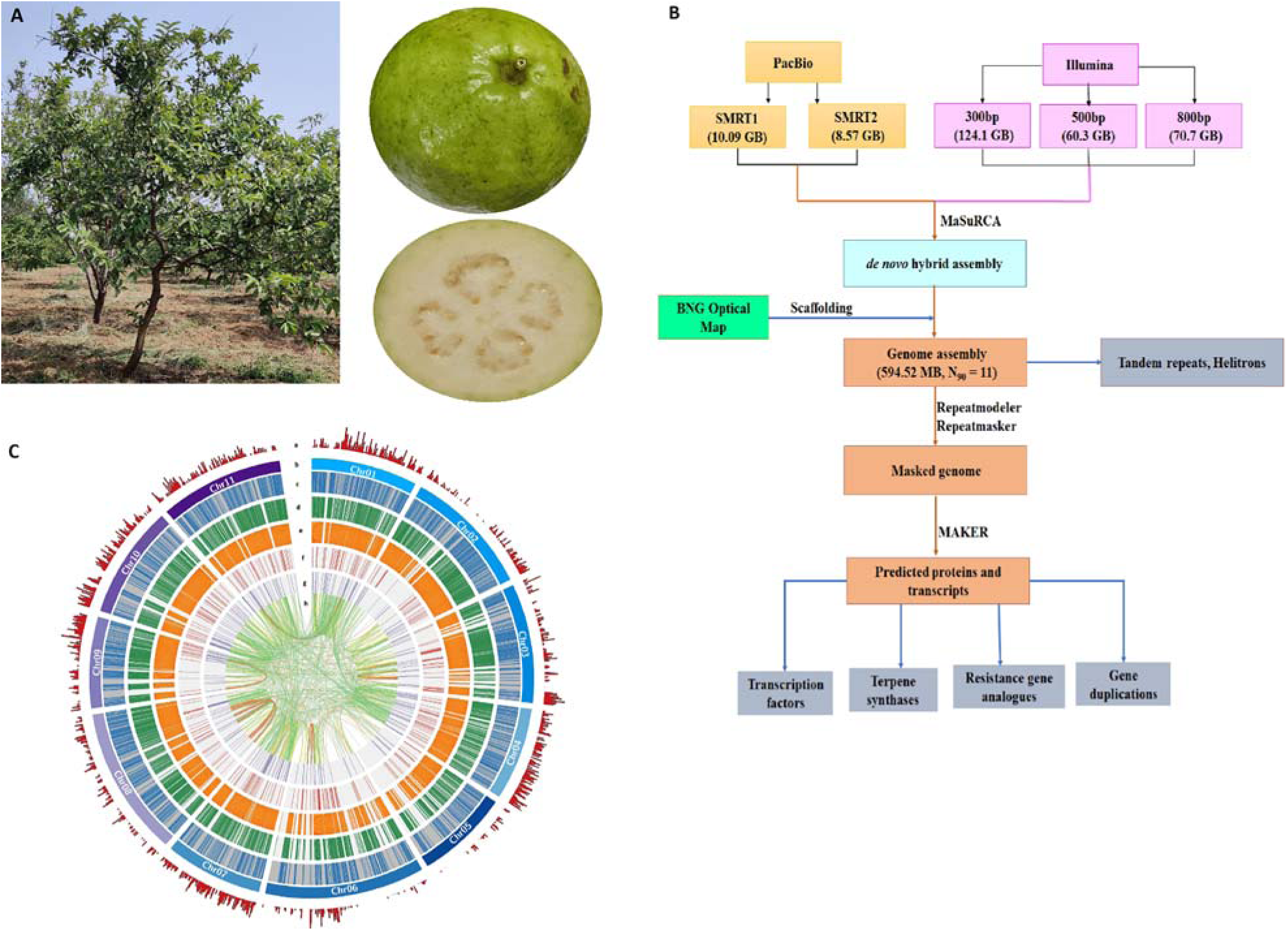
Guava cv. Allahabad Safeda genome sequencing, assembly, and annotation. **A.** Ten-year-old tree and fruit **B.** Schema of genome sequencing, assembly, and annotation **C.** Circos plot representation of a. gene density b. Pseudochromosomes c. LTR repeats d. Tandem repeats e. LINES, SINES & Helitrons f. Non-coding RNAs g. Transcription factors h. Duplications.

Three paired end libraries of 300, 500 and 800 bp insert size were constructed and 827.935470 M, 402.367298 M, 471.789730 M reads were generated, respectively with Illumina sequencing, leading to a total of 255.31388 GB short read data (Figure 1B, Table S1). Two independent SMRT cells of PacBio Sequel generated 10.09 and 8.57 GB polymerase read bases with 788,685 and 794,336 reads with mean polymerase read length of 12.804 and 10.794 Kb. Polymerase read N50 from 2 SMRT cells was 20,649 and 17,664 bp. Adapter-removed (Cutadapt 1.8) reads were subjected to *de novo* hybrid assembly with MaSuRCA(Zimin et al., 2013) ver. 3.3.1. DLE-1 enzyme restriction of HMW AS genomic DNA followed by sequencing allowed 451.4270 GB sequence data for BioNano maps. The hybrid assembly scaffolded with BioNano optical genome map generated a final scaffold sequence assembly of 594.521 MB and contig sequence of 479.656 MB. The genome scaffold N90 is 11, suggesting a chromosome scale assembly. Benchmarking Universal Single-Copy Orthologs (BUSCO) analysis for 2121 eudicot genes (Figure S1), Quality ASsessment Tool (QUAST) evaluation (Figure S2) and LTR assembly index (Figure S3) further indicated a high-quality chromosomal level genome assembly. A GenomeScope K-mer frequency distribution plot (Figure S4) indicated 0.15% heterozygosity in the AS guava genome. The largest scaffold is 62.699 MB long representing the largest guava chromosome.

With AS transcriptome(Mittal et al., 2020) as evidence we predicted 17,395 genes with MAKER (Cantarel et al., 2008) (Supplementary Datafile1; that are 23% more than the draft assembly (Thakur et al., 2021)). Comparison of all 17,395 predicted genes’ annotation against eudicots (OmicsBox v.3.0.27) with Blast2GO identified gene ontology for 12,613 genes. InterProScan and PfamScan found domains for 16,302 and 13,774 genes (Table 1, Supplementary Datafile2). The BRAKER pipeline predicted 59,448 genes in the assembly and 42,428 could be functionally annotated with Uniprot database using BLASTX program with E-value cutoff of 10^-3^ However, MAKER pipeline output for guava genome characterisation was used owing to better average gene length (data not shown).

**Table 1:** Genome assembly statistics of reference and comparison to the draft.

| Genome assembly | Draft<br>(Number / Size) | Reference<br>(Number / Size) |
| --- | --- | --- |
| Scaffolds | 45455 | 2322 |
| Scaffold N50 | 17.992 KB | 50.573 MB |
| Scaffold N90 | 3.312 KB | 39.769 MB |
| Total gene length | 62.43917 MB | 84.310715 MB |
| Average gene length | 4.423373503 KB | 4.8468361598 KB |
| Longest Scaffold | 134.502 KB | 62.699 MB |
| Assembly Length (MB) | 303.782 | 594.521 |
| GC content (%) | 39.51 | 39.85 |
| BUSCO | 89.9% | 95.9% |
| Complete and single copy | 87.3% | 92.1% |
| Complete and duplicated | 2.6% | 3.8% |
| Missing | 4.3% | 3.4% |
| Fragmented | 5.8% | 0.7% |
| **Genome Annotation** |  |  |
| Repeat content (%) | 37.95 | 39.39 |
| Predicted genes | 14115 | 17395 |

| Gene Annotation |  |  |
| --- | --- | --- |
| InterProScan | 11686 | 16302 |
| Pfam | 10981 | 13774 |
| Blast2GO | 11979 | 12613 |

The genomic features of reference genome assembly *viz.* gene density, pseudochromosomes, LTR repeats, tandem repeats, LINES, SINES & Helitrons, non-coding RNAs, transcription factors and duplications were identified (Figure 1C). We identified 647 tRNAs with tRNASCAN-SE out of which 573 code for standard amino acids, 4 with unknown isotypes, 24 retained introns and 70 were putative tRNA pseudogenes (Supplementary Datafile3). StructRNAfinder predicted 701 miRNA, 917 rRNA, 100 snRNA and 313 snoRNAs (Figure 1C, Supplementary Datafile4). Krait(Du et al., 2018) identified 125984 perfect microsatellites covering 2534106 bp with relative density of 5283.17 bp/Mb (Figure S5) with 1435 found in CDS region, 4768 in exonic region and, 17331 in intronic region. Compound SSRs, Imperfect SSRs (iSSRs) and Variable SSRs (VNTRs) were 5000, 618550 and 46945 at relative densities of 574.86 (bp/Mb), 38295.71 (bp/Mb) and 1839.05 (bp/Mb).

### Classification of guava retroelements

Approximately 39.39% of the genome was repetitive with 25.63% consensus repeats and 13.76% unclassified unique to guava AS (Table S2). LTR elements were dominant repeat types constituting 16.74 % of the genome length with Gypsy and Copia 9.1% and 7.31%, respectively (Figure 2A; Table S2; Supplementary Datafile5). DNA elements constituted 6.69% with 2.27%

**Figure 2.**
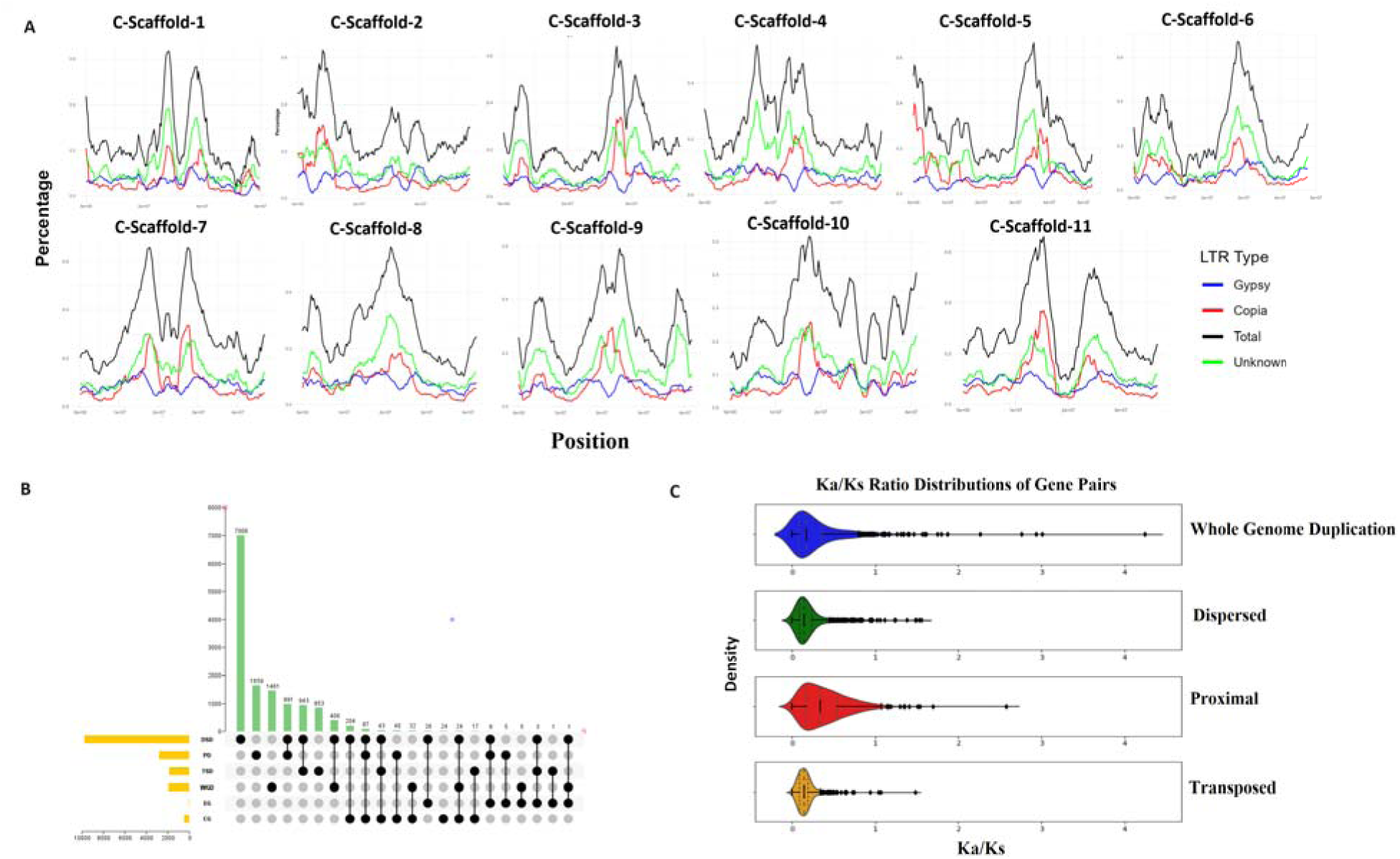
Guava cv. Allahabad Safeda genome characterization. **A.** Chromosome wise LTR distribution of Gypsy, Copia and unknown function **B.** Upset plot depicts the overlap of major genomic duplications **C.** Violin plot represents Ka/Ks distribution of duplicated gene pairs

MULE-MuDR elements. Non-LTR retrotransposon, LINEs (1.3%) and SINEs (0.02%), accounted for a small proportion of the total repeats. More than 62,000 tandem repeats (Supplementary Datafile6) and 886 Helitrons (Supplementary Datafile7) were present in the genome. The highest Helitron density was found on Chr10 (Supplementary Datafile8). LTR_retriever found 7722 undamaged LTR-RTs in guava. The intact LTR C-Scaffold-2:40085060..40087627_LTR/Gypsy had the highest copy number 1559 (0.57424%) among Gypsy (Figure S6) and LTR C-Scaffold-1:43010129..43010347_LTR/Copia copy 1500 (0.06046%) among Copia elements (Figure S7 & Supplementary Datafile9). Sub-classification of LTR-reterotransposons identified Copia/Ale and Gypsy/Tekay as predominant while Copia/Angela/Alesia and Gypsy/Chlamyvir as least abundant classes (Supplementary Datafile10). The distributions of insertion times showed that Gypsy and Copia in guava appeared recently with increased insertion activity from 0 to 5 MYA (Figure S8A & S8B) and found phylogenetic similarity among intact (containing GAG, PROT, RH, RT, INT domains) LTRs (Figure S8C). Many attributes pertaining to processes such as development, biosynthesis and pest resistance might potentially be shaped by these retroelements and would be interesting subject for revelation.

### Gene duplications, gene family expansion/contraction, transcription factors, terpene synthases, resistance genes and guava speciation

Gene duplication is thought to be a fundamental driver of genome evolution, allowing organisms to adapt to environmental changes(Ames et al., 2010; Magadum et al., 2013; Zhang et al., 2020). Gene duplication detection with DupGen_finder (Table S3) suggested expansion of most genes was dominated by Dispersed Duplications (DSD-58.39%; Figure 2B). Approximately, 12.5 % of gene pairs were retained from transposed - TRD followed by proximal - PD (10.14%) and whole genome -WGD (5.69%) duplications. Ka/Ka and Ks distribution analysis for gene duplications originating through dispersed and proximal duplicate gene pairs showed lower Ks values (Figure S9) and, higher Ka/Ks ratios (Figure 2C & Supplementary Datafile11) suggestive of an ongoing, continuous sequence divergence and stronger positive selection. In brief, newly generated dispersed duplicates (DD) and proximal duplications (PD) contributed to gene family contraction and expansion in *Psidium guajava*.

Our comparative analysis of the guava genome to 11 other angiosperms including *E. grandis*, *S. grande*, and *M. polymorpha* from Myrtacea, revealed 2056 unique genes exclusive to guava and a higher degree of overlap among members of Myrtacea (Figure S10) than among more distant taxa. Gene ontology (GO) enrichment analysis of 282 single copy clusters revealed association with a wide range of significant biological, molecular, and cellular functions (Figure S11 and Supplementary Datafile12). Guava has undergone both gene family expansions (17) and contractions (388), albeit to a lesser extent than other species within the Myrtaceae family (Figure S12 and Supplementary Datafile13). GO enrichment analysis (Figure S13 & Supplementary Datafile14, 15) revealed expansion of gene families for cellular response to amino acid stimulus and monoterpene biosynthetic process, and contraction of gene families involved in response to salt stress and water deprivation.

Within the phylostratum *’Psidium guajava*,’ 524 genes were found orphan / unique to guava. Phylogenetic analysis based on single-copy clusters confirmed the placement of *P. guajava*, *E. grandis*, *M. polymorpha* and *S. grande* within the Myrtaceae family in a monophyletic group (Figure S12 and Figure S14). Phylostratigraphic analysis found guava genes distributed across phylostrata representing evolutionary stages of development (Supplementary Datafile16). Although 239 genes were not enriched for ontologies related to any biological activity, 285 orphan genes were associated with diverse functional roles including important enzyme-, transporter-, stress response- and defense-related gene ontologies.

Transcription factors (TFs) and regulators (TRs) are key for understanding of fundamental mechanisms related to environmental adaptation and plant development(Dai et al., 2013). Web-based tool iTAK identified 875 TFs (Table S4) and 325 TRs (Table S5) in the AS genome. Among the 63 TF classes, MYB (152), WRKY (51) and bHLH (74) were over-represented in the guava genome with highest representation of the functionally diverse and most abundant MYB family (Peng et al., 2016). Independent homology-based prediction identified 69 MYBs (Supplementary Datafile17) with specific domains - 15 R2R3, 53 1R and a single R3. Phylogenetic analysis of MYBs among *A. thaliana*, *M. domestica*, *V. vinifera*, *F. vesca*, *P. persica* and guava classified them into four clades of secondary metabolites *viz.* Proanthocyanin, anthocyanin, flavanols and flavonoids (Figure 3A). Guava MYB proteins mostly with clustered together, but also exhibited significant similarity to MYBs of other species. Among 23 TR classes, SNF2, playing roles in plant development, DNA repair and transcription regulation(Guo et al., 2022), was the most represented followed by PHD TR with role in histone binding, pathogen defense responses and plant development (Mouriz et al., 2015).

**Figure 3.**
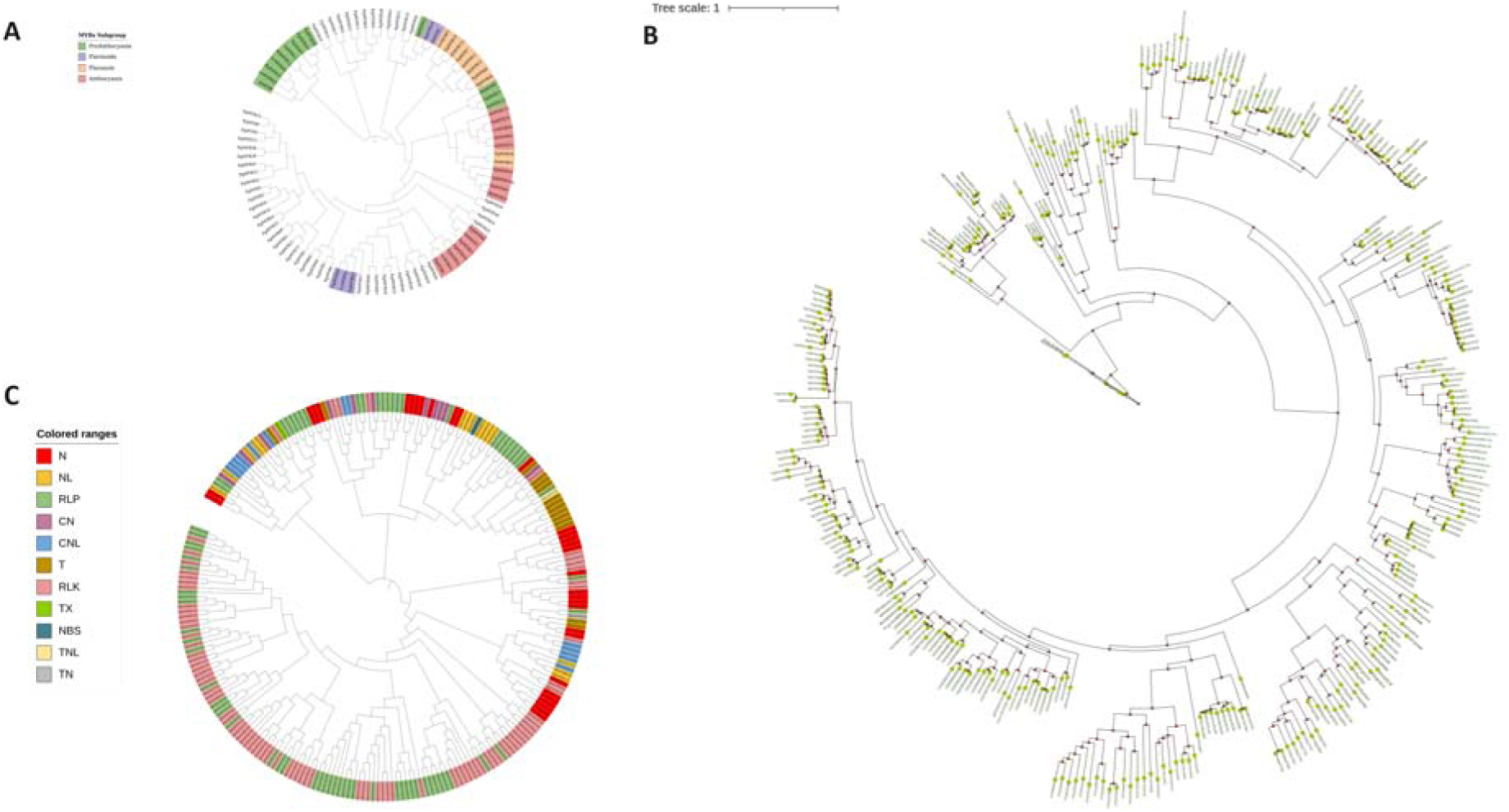
Phylogenetic tree of. **A.** MYB transcription factors of *Arabidopsis thaliana*, *Malus domestica*, *Vitis vinifera*, *Fragaria vesca* and Prunus persica **B.** Terpene synthases of *Malus domestica*, *Prunus persica*, *Arabidopsis thaliana*, *Populus trichocarpa*, *Vitis vinifera*, *Sorghum bicolor*, *Oryza sativa*, *Selaginella moellendorffii*, and *Physcomitrium patens*. **C.** Resistance gene analogues of guava identified RGAugury, RRGpredictor, DRAGO2 and NLR-annotator2.

Terpenes / terpenoids are important naturally occurring aromatic compounds in plants with biological roles in attraction of pollinators, fragrance and pigmentation of leaves, flowers, fruits, seeds, roots (Chemical Constituents of Grapes and Wine, 2008) and as major constituents of essential oils (Cox-Georgian et al., 2019). Aroma of guava fruit and leaves is reminiscent of resinous fragrance from eucalyptus and Indian Black berry that are rich in terpenes. The TPS gene family has eight different subfamilies categorised by the structurally distinct terpenoid compounds they synthesize, viz. TPS-a (sesqui-terpene), TPS-b and TPS-g (cyclic/acyclic mono-terpene), TPS-c and TPS-e_f (copalyl diphosphate, ent-kaurene, and di-, mono- and sesqui-terpene). Subfamilies TPSd and TPSh synthesize gymnosperm and *Selaginella* specific terpenes respectively. In AS, 47 TPS genes were identified using Terzyme (Supplementary Datafile18) and functionally classified into monoterpene (23), diterpene (22), and sesquiterpene synthases (2). The TPSb gene family is found most represented followed by TPSe_f and TPSg (Table S6). Eucalyptus, also from the Myrtacea family, had the highest number of TPSa gene(Külheim et al., 2015), which were least abundant in guava. However, TPSe_f and TPSg copy number are similar in eucalyptus and guava. Interestingly, no TPSh gene family was found in guava. Phylogenetic analysis of TPS protein sequences of guava placed TPSb, TPSg and TPSe_f in their respective clades with few exceptions (Figure S15a). Further, homology search for TPS in the N-terminal domain - PF01397 and metal binding domain - PF03936 found 33 full length terpene synthases in guava genome (Supplementary Datafile19). Guava TPS phylogeny with 9 plants/ lycophytes (IQ-TREE - best-fit model - JTT+F+G4; Figure 3B) shows close relationship with eucalyptus TPS genes. Expansion of monoterpenoid biosynthetic process GO:0016099 (∼20% expanded gene families – Supplementary Datafile14) and high transcript abundance of TPSb in leaves (Figure S15B) vouches for extra dosage of monoterpenes in guava.

To explore the defence system of guava we predicted resistance gene analogues (RGAs) (Supplementary Datafile20) with RGAugury, RRGpredictor, DRAGO2 and NLR-annotator. We found NBS, LRR, TIR and CC domain classes of resistance genes indicating a robust defence system against diseases. Guava proteins interrogated with RGAugury found 146, RRGpredictor identified 367 and DRAGO2 identified 726 RGAs in guava belonging to TM-CC, CN, CNL, NL, RLK, TX, N, RLP, TN, TNL, L, KIN and CK classes. Out of all prediction tools DRAGO2 had highest sensitivity and retrieved most of the classes. Overall, TM-CC, RLK, RLP and KIN were the most abundant classes while TN and TNL were the least frequent classes in guava. RGAs in guava are unevenly distributed with maximum on chromosome 4 followed by chromosome 7 (Supplementary datafile21) suggesting key role of these chromosomes in defence mechanisms. NLR-annotator identified 785 resistance genes (Supplementary datafile22) on guava genome where the TIR-NLR domain combination was most frequent followed by CN, NL, CNL, TN and TX. Phylogenetic analysis resistance gene sequences groups in several clades, notably RLK and RLP, found significant homology among RGA classes (Figure 3C). This finding suggests that these two classes of RGAs are evolutionarily conserved and may have similar functions across different plant species. Overall, the identification of RGAs in guava provides insights into the defence mechanisms of this fruit tree and may help in the development of disease-resistant varieties.

### Guava underwent a single polyploidy event like other Myrtales

Colinearity/synteny patterns between *Psidium guajava*, *Eucalyptus grandis*, *Syzygium grande*, *Metrosideros polymorpha*, *C. citiodora* and *Rhodamina argentea* supported previous studies finding at least one whole genome duplication event and a shared polyploid past (Figure S16). Comparing the genomes of *Populus trichocarpa* and *V. vinifera* with that of guava, clear 2:1 syntenic patterns indicate independent whole-genome duplication (WGD) specific to the Salicaceae and Vitaceae family (Figure S17). Synonymous substitution (Ks) values between orthologues of guava and other Myrtacea [(*E. grandis* (0.19926452), *S.grande* (0.18499773), *M. polymorpha* (0.15675545), *R. argentea* (0.10627856), *C. citiodora* (0.19209344)] are much lower than between guava compared to *P. granatum* (1.20703292) and *V. vinifera* (1.29041193) of family Lythraceae and Vitacea, suggesting a Myrtacea-wide evolutionary event (Figure S18).

### Lycopene in Pink pulp of guava is controlled by Phytoene Synthase 2 and is associated with diagnostic genomic structural variation

Lycopene estimation of 20 genotypes revealed that white-pulp types had non-significant amount of lycopene (0.019 – 0.6 mg /100 gm pulp) while the pink- and yellow types had a range of 1.8-6.2 mg/100gm. Candidate gene-based screening of 47 carotenoid biosynthesis genes and/or different isoforms (Table S7; Figure 4A) with Illumina based genome re-sequencing of 20 genotypes and data visualization with integrative genome viewer(Robinson et al., 2011) revealed major structural variations on pseudochromosome 5 (26201203 – 26204356) in pgpauas08401. It is a Phytoene Synthase (*PSY*2) gene, the first committed step in lycopene biosynthesis pathway. *PSY*2 in guava is 3153 nucleotide long gene with 5′-UTR, 4 exons, 3 introns and 3′-UTR. The gene had 3 exonic and 6 intronic SNPs (Table S8, Figure 4B) and a 5/6 AG dinucleotide repeat (10-12bp) deletion 17bp downstream of 3′- UTR in pink pulp genotypes. A primer pair designed (Table S9) from the region flanking the deletion amplified -100 bp DNA amplicon in each of 12 white fleshed types, and two bands -90/88 and -100 bp in each of eight pink fleshed genotypes (Figure 4B).

**Figure 4.**
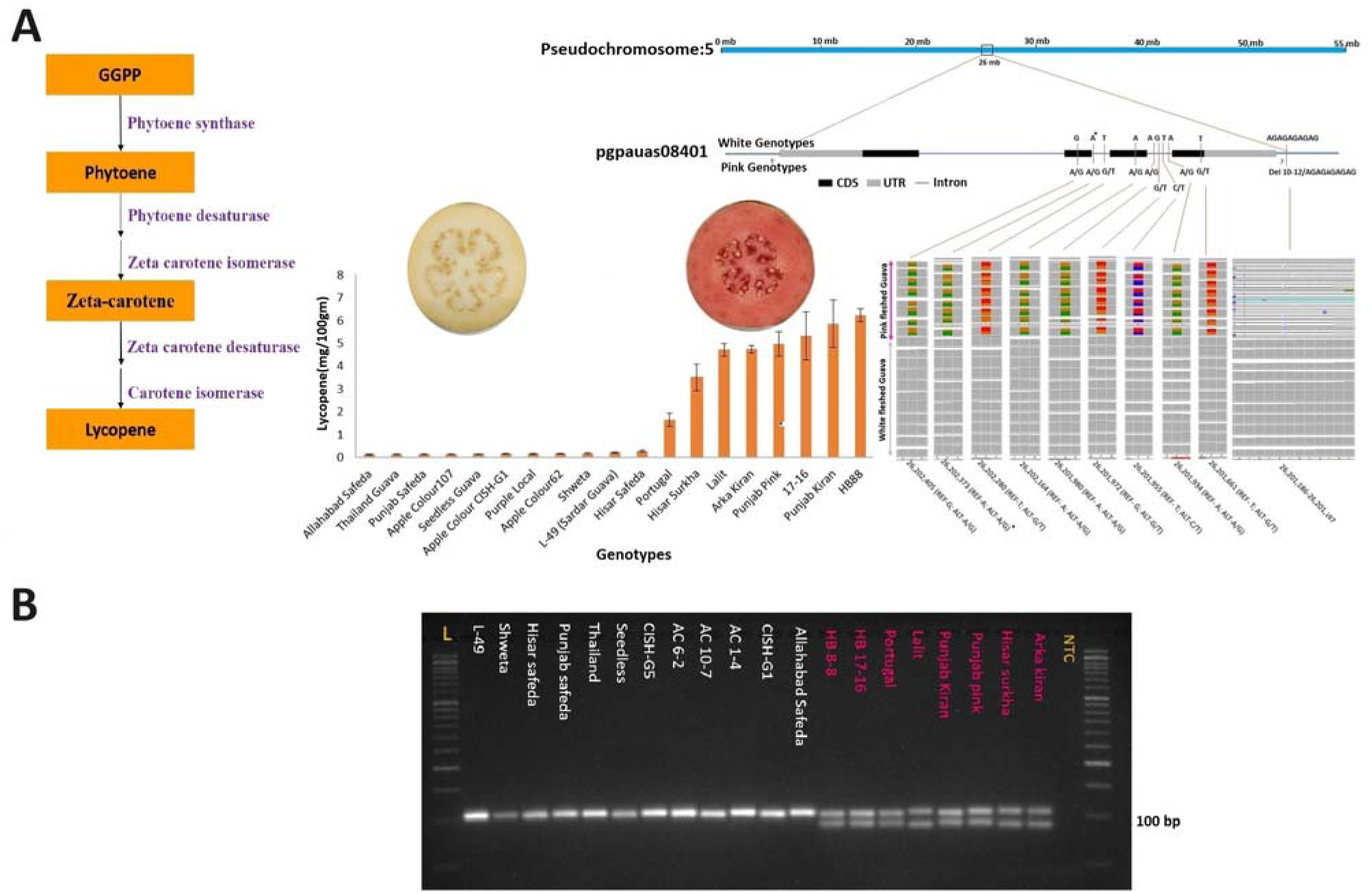
Candidate gene for pink pulp colour, lycopene content variation and structural genetic variability. **A.** Minimal pathway for lycopene synthesis, gene structure of Phytoene Synthas 2 (pgpauas08401) in white fleshed guava cv. Allahabad Safeda, *in-silico* structural variations visualized by re-sequencing data in 20 pink and white fleshed genotypes on integrated genome viewer and lycopene variability. **B.** The dinucleotide repeat (AG)_5/6_ deletion 17bp downstream of 3′-UTR in pink genotypes in heterozygous condition leads to dual bands of the diagnostic marker visualised on 3.5% agarose gel for pink pulp.

Lycopene estimation over a large panel of 82 guava cultivars/accessions showed similar results, with 67 white genotypes having non-significant amount of lycopene and the -100 bp DNA amplicon, while the 15 yellow- and pink-coloured genotypes had lycopene ranging from 1.2-7.6 mg/100gm and the two bands -90/88 and -100 bp (Figure 5A-B). Further, we performed double digest restriction-site associated DNA sequencing of the 82 genotypes and genome wide association with GAPIT3(Wang and Zhang, 2021) using CMLM and FarmCPU model, revealing strong association of pink pulp colour trait on Psuedochromosome 5 (Table S10, Figure 5C). The causal A/G SNP identified by ddRAD is one of the SNPs found earlier by genome re-sequencing, suggesting a strong association to pink pulp trait. Another strong association is found on Psuedochromosome 5 in reticulon like protein B8 isoform X1 (pgpauas08392) 120,987 bp upstream of pgpauas08401. Reticulons are endoplasmic reticulum (ER) membrane associated proteins that help in ER shaping. However, understanding the reticulon association to pink pulp of guava is still an open question.

**Figure 5.**
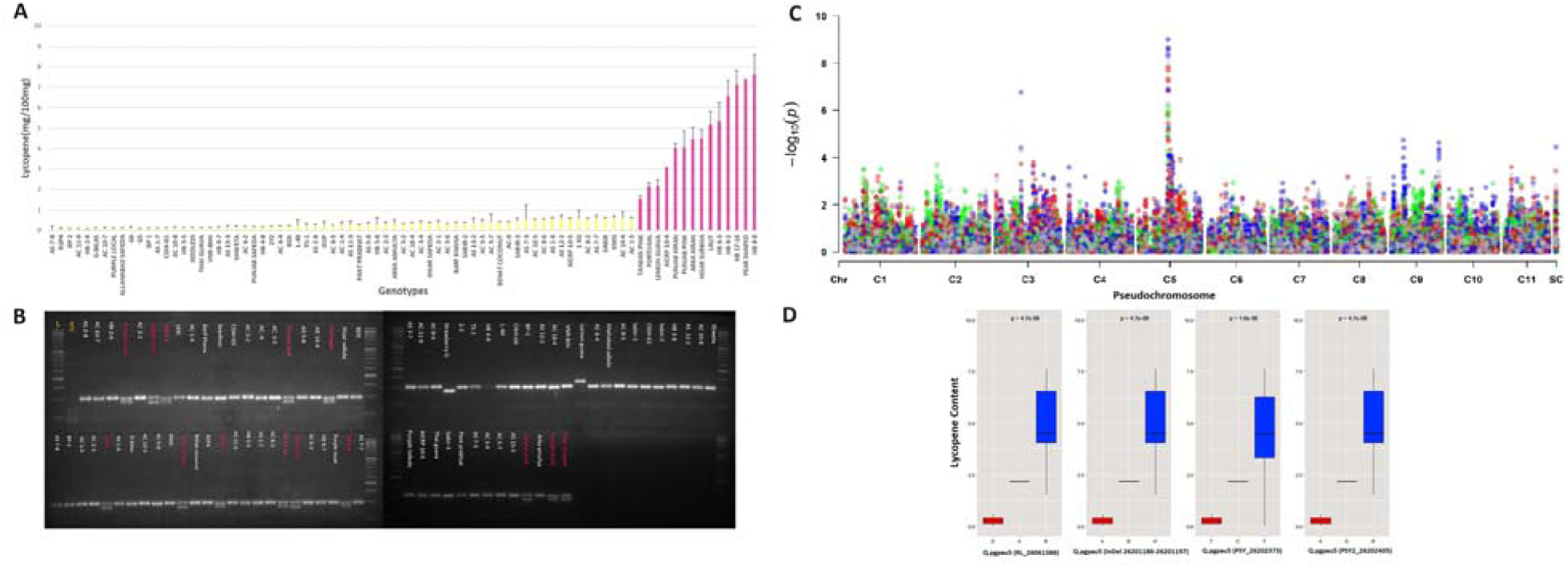
Lycopene content variation in Indian guava germplasm core-set and association to Phytoene Synthase 2 (pgpauas08401) **A.** Lycopene content variation in 67 white fleshed and 15 pink fleshed genotypes. **B.** Diagnostic deletion in Phytoene Synthase 2 (pgpauas08401) is associated in all the pink genotypes in heterozygous condition leading to dual band appearance on 3.5% agarose gel compared to single band of 100bp in white fleshed genotypes **C.** Genotyping by sequencing with ddRAD on 82 genotypes with 67 white- and 15 pink-fleshed genotypes identify strong association on chromosome 5 harbouring Phytoene Synthase 2 (pgpauas08401) and reticulon like protein B8 isoform X1 (pgpauas08392) 120,987 bp upstream of pgpauas08401 visualised by Manhattan plot. **D**. Box plot of lycopene content vs allelic variation among pink vs white guava genotypes (Blue boxes – pink pulp genotypes; Red boxes – white genotypes)

*In silico* protein interactome analysis with STRING established the interactomes of PSY2 (Figure S19) evidencing 11 putative protein interactors of order ∼0.9 (Table S11) along most relevant enzymes with established roles in lycopene biosynthesis like 15-cis-phytoene desaturase (PDY) and geranylgeranyl pyrophosphate synthase (GGPP).

## Discussion

A high-quality reference genome expedites discovery and utilization of the full genetic potential of a species, to cater the needs of human race. Advances in NGS technologies has made it easier to sequence, assemble, annotate large plant genomes. Although many genomes have been sequenced using doubled haploid individuals in fruit species like grape and potato, short read sequences for heterozygous species remained difficult to assemble until recent computational algorithms advancements. Our interventions for generating doubled haploid in guava remained futile(Sohal et al., 2019) so we proceeded to generate chromosomal level assembly of a heterozygous individual of Allahabad Safeda using refined computational algorithms. This guava genome assembly with a scaffold N50 of ∼50.5 Mb is higher than the previously reported New Age (NA) guava genome(Feng et al., 2021) (N50 = ∼40 Mb).

Synteny of AS to NA genome (Figure S7) found a correlation of ∼ 0.89 with only a few genome rearrangements like inversion events indicating minor evolutionary divergence leading to big changes such as high (∼550 mg/100g fresh weight) vitamin C content(Feng et al., 2021) in mature NA fruit, similar to the levels in Indian gooseberry which is much higher than ∼200 mg/100g fruit weight as in Indian guavas like that of AS.

Identification of terpene synthase, resistance genes and transcription factors vouch for the high quality of the assembly. Terpene synthases are highly expressed in guava leaf tissue, similar to eucalyptus(Külheim et al., 2015) whereas high content in flower buds / open flowers followed by young tendrils and leaves(Matarese et al., 2014) is found in grapes. Expansion of monoterpene synthases gene family in guava demands quantifying monoterpenes in guava tissues for identifying the natural source of it to be utilised as bio-pesticides.

The gene number in guava genome is lower than expected, so a next step forward should be developing a pan-genome for identifying the missing genes. Running iso-seq and generating full length transcripts for gene evidence will aid in developing improved gene models. Development of F_1_ populations of pink fleshed x white fleshed cultivars and the progeny thereof segregating for fruit colour and its potential association with a PSY2 gene marker will mark the utilization of this genome assembly in enhancing the guava breeding program by reducing acreage, time, and labor for selecting pink pulp segregants at seedling stage. Also, the genome assembly developed here will aid in chromosome walking for high resolution mapping of other traits of commercial importance like metabolites of nutraceutical importance in peel and pulp, seed strength and/ number, fruit core size, total sugars and many more.

## Methods and Materials

### Plant Material, nucleic acids extraction and sequencing

Guava germplasm is maintained at Fruit Research Farm, PAU, Ludhiana and at Fruit-research Station, Bahadurgarh, Patiala, Punjab. Guava cv. Allahabad Safeda (AS) etiolated leaf samples were subjected for high molecular weight (HMW) DNA extraction using ‘IrysPrep Plant Tissue DNA isolation kit’ (young leaves of AS tree growing in orchard covered with black chart for ∼10 days for etiolation). The samples were run on Pulse Field Gel electrophoresis system and DNA sample with average molecular weight of ∼200-150 kb was selected for sequencing on PacBio Sequel and Saphyre for Bionano maps. Libraries were developed using ‘SMRTbell template preparation kit’ following the manufacturer’s instructions and sequenced on the PacBio sequel platform (https://www.pacb.com). BioNano optical maps were developed with BioNano Genomics (https://bionanogenomics.com) protocols. About 1µg HMW DNA was used for DLS labelling using ‘DNA labeling kit-DLS’ and labelled DNA was imaged on the Saphyr system (https://bionanogenomics.com) in a lane of Saphyr Chip. RefAligner was used to identify all molecule overlaps, and a consensus map was constructed. The BioNano IrysSolve module ‘HybridScaffold’ was subsequently used for generating hybrid assembly between PacBio-SMRT contigs and BioNano-assembled genome maps. Completeness and continuity of the final assembly was estimated using various methods BUSCO(Waterhouse et al., 2018), GenomeScope(Vurture et al., 2017), QUAST(Gurevich et al., 2013) and LAI(Ou et al., 2018).

DNA extraction of 12 white fleshed genotypes *viz.* L-49/Sardar guava, Arka Amulya, Shweta, Punjab Safeda, Hisar Safeda, Seedless, VNR-Bihi, CISH-G5 (Lalima), AC 1-4, AC 6-2 (Punjab Apple Guava), AC-10-7, CISH-G1 and 8 pink/yellow fleshed genotypes Lalit, Arka Kiran, Punjab Pink, Punjab Kiran, Hisar Surkha, HB-88, HB-17-16, Portugal was performed with Qiagen DNeasy^®^ Plant Mini Kit to be used for genome re-sequencing. Illumina libraries of insert size 800 bp, 500bp and 300 bp were prepared for AS and 300 bp for all other genotypes following manufacturer’s protocol (https://illumina.com). The PE libraries were sequenced on Illumina HiSeq X Ten sequencer.

### Annotation and characterization of Chromosomal level Allahabad Safeda assembly

The *de novo* repeat identification was performed with RepeatModeler v 1.0.1. RepeatMasker-version 2.1 (http://www.repeatmasker.org) for complete identification of repetitive sequences as well as for masking of the genome, and the masked genome was further used for annotation. Krait(Du et al., 2018) software was used for identification of microsatellite and their distribution over the genome. SINEs and Helitrons were also identified with SINEfinder and EAhelitron(Hu et al., 2019) with default parameters. Tandem Repeat sequences prediction was done with TRF (Tandem Repeats Finder(Benson, 1999)) algorithm with default parameters. LTR retrotransposons predictions was done using two separate pipelines *viz.* the LTRharvest(Ellinghaus et al., 2008) & LTRdigest pipeline(Steinbiss et al., 2009) and LTR-FINDER(Xu and Wang, 2007). The LTR candidate predictions from LTRharvest and LTRFinder were fed to LTR_retrivar(Ou and Jiang, 2018) for predicting whole genome LTR-RT annotation. Search for gypsy and copia was done as if RNase H, reverse transcriptase, and integrase domains were present. Order of these 3 domains was used to classify Ty1/Copia (integrase upstream of RNase H) and Ty3/gypsy (RNase upstream of integrase) superfamilies. The sub-classification of gypsy and copia was done using TESorter(Zhang et al., 2022) using REXdb(Neumann et al., 2019) and GyDB(Llorens et al., 2011), Alignment of the intact LTRs was done using MAFFT(Nakamura et al., 2018) and Phylogenetic tree was constructed using IQ-TREE(Nguyen et al., 2015) and plotted using iTOL(Letunic and Bork, 2021). Insertion time calculation was done using the pipeline included in LTR_retrivar.

Transfer RNA (tRNA) genes were predicted using tRNAscan-SE version 2.0.5(Chan et al., 2021) using default parameters and for prediction of other classes of RNA - StructRNAfinder(Arias-Carrasco et al., 2018) was used. The MAKER v 2.31.10 pipeline(Cantarel et al., 2008) was used for prediction of the genes in the masked genome assembly. AS transcriptome assembly(Mittal et al., 2020) was used as EST evidence. To generate *ab initio* gene predictions with the masked assembly SNAP(Korf, 2004) and AUGUSTUS v 3.2.2(Stanke et al., 2006) were used. MAKER was run for three consecutive rounds for prediction of genes as described before(Thakur et al., 2021).

All the predicted genes were annotated by comparing the protein sequences with PfamScan (http://www.ebi.ac.uk/Tools/pfa/pfamscan). Protein domain identification was done with InterProScan version 5.39-77.0 (http://www.ebi.ac.uk/interpro/download/). The transcript sequences of predicted genes were subjected to BLAST against Viridiplantae in OmicsBox v.3.0.27 and annotated using plugged-in Blast2GO.

Terpene synthase (TPS) genes were identified with Terzyme(Priya et al., 2018). Multiple alignments of all TPS protein sequences for phylogenetic analysis was done with CLUSTAL-W ver.2.1(Larkin et al., 2007) and the tree was constructed with iTOL(Letunic and Bork, 2021). Terpene genes were further predicted by manually searching guava proteins for HMM profiles using InterProscan, HMMScan and PFamScan. Any protein containing PF01397 (N-terminal TPS domain) or PF03936 (metal binding domain) or both were identified as TPS genes. Terpene synthases genes from *Malus domestica*, *Prunus persica*, *Arabidopsis thaliana*, *Populus trichocarpa*, *Vitis vinifera*, monocots: *Sorghum bicolor*, *Oryza sativa*, lycophyte: *Selaginella moellendorffii* and moss: *Physcomitrium patens* were retrieved. Amino acid alignments were made with MUSCLE module of Genious prime using standard parameters. To generate phylogeny, we first tested for amino acid substitution model providing the maximum likelihood tree with best Akaike’s information criterion (AICc) value. The tree with highest AICc value was selected and phylogeny was determined using 1000 bootstrap replicas followed by construction with iTOL.

TFs were identified in the protein sequences with iTAK(Zheng et al., 2016) & PlantRegMap(Tian et al., 2020) and TRs with iTAK. A homology-based search was also done using TFs to identify the potential members of MYB TF gene families in guava genome. Hidden Markov Model (HMM) profiles of MYB (PF000249) were downloaded from Pfam and used as the query to search against protein sequences databases employing hybrid approach of using Pfamscan, InterProScan and HMMscan, with E-value thresholds set to 1e^-10^. Outputs of iTAK and Arabidopsis HMM profiles were combined and redundancy was removed with TBtools(Chen et al., 2020). Identified protein sequences were manually inspected with SMART (http://smart.embl-heidelberg.de/) to verify the presence of conserved domains. MYB protein sequences of *A. thaliana* were downloaded from TAIR, and sequences of *M. Domestica*, *V. Vinifera*, *F. vesca* and *P. persica* from phytozome. Subsequently CLUSTALW2 was used for multiple sequence alignment and phylogenetic tree was constructed with iTOL. The protein properties like isoelectric point and molecular weight of the predicted TF were calculated using ProtParam server’s wrapper script (GitHub - zmactep/ProtParam.jl).

RGAugury(Li et al., 2016), RRGpredictor(Santana Silva and Micheli, 2020) and DRAGO2(Osuna-Cruz et al., 2018) were used to identify different classes of resistance gene analogues (RGAs). NLR-annotator2(Steuernagel et al., 2020) was used for identification and validation of NBS-LRR. The resistance genes were subjected to multiple sequence alignment with CLUSTALW2, and phylogenetic tree was constructed with help of iTOL.

### Genome duplications and Whole genome Synteny Analysis

We investigated the frequency of genome-wide duplications in guava. DupGen_finder(Qiao et al., 2019) pipeline was used for identifying whole-genome duplicates (WGD), tandem duplicates (TD), proximal duplicates (PD), transposed duplicates (TRD), and dispersed duplicates (DSD).

Using the KaKs_Calculator (version 3.0) (Zhang, 2022), we further determined the Ka (number of substitutions per nonsynonymous site), Ks (number of substitutions per synonymous site), and Ka/Ks values for gene pairs after conversion to codon alignment using PAL2NAL(Suyama et al., 2006) and produced by the various duplication modalities based on the YN model. Genome assembly and annotation for *Psidium guajava* were uploaded to the CoGe (Lyons and Freeling, 2008) comparative genomics platform online. With CodeML configured to “Calculate syntenic CDS pairs and color dots: Synonymous (Ks) substitution rates,” Ks calculations were performed to create syntenic dot plots. Plant species with CoGe genome IDs *Psidium guajava* (id65865), *Eucalyptus grandis* (id28624), *Punica granatum* (id41383), *Populus trichocarpa* (id38424), *Metrosideros polymorpha* (id45020), *Syzygium grande* (id42510) and *Vitis vinifera* (id30669) using Quota Align syntenic depth of 2:1, max query chromosomes = 100, max target chromosomes = 25, and “Use all genes in target genome” were used for each pairwise SynMap(Haug-Baltzell et al., 2017) analysis (including self:self). The mapping of Populus against Psidium employed a Quota Align(Tang et al., 2011) syntenic depth of 2:2 and the same settings as for depth 2:1. Ks values for syntenic paralogs were plotted as density plots.

### Ortholog, gene family and phylostratigraphy analysis

We conducted a comparative genomic analysis of eleven plant species *Eucalyptus grandis*, *Gossypium raimondii*, *Brassica rapa*, *Populus trichocarpa*, *Malus domestica*, *Vitis vinifera*, *Punicia granatum*, *Metrosideros polymorpha, Citrus clementina*, and *Syzygium grande* with *Psidium guajava*. The protein sequences of these species were retrieved from publicly available databases, including GenBank and the respective genome sequencing projects. OrthoMCL(Li et al., 2003) was deployed to identify orthologous gene relationships among the species. The resulting orthologous gene clusters were then utilized to investigate conserved genes, gene family expansions or contractions, and potential gain or loss of genes across the plant species using CAFE5(Mendes et al., 2020). To investigate the evolutionary background of guava, single-copy gene clusters were treated as independent evolutionary units. MUSCLE was used to align protein sequences, trimAl(Capella-Gutiérrez et al., 2009) to remove and trim conserved sequences, and FastTree(Price et al., 2009) to construct the phylogenetic tree between species using the maximum likelihood technique. Each phylogenetic node’s veracity is evaluated using the SH test procedure. The divergence time estimation was done by the GTR substitution model, using the software MCMCTree from PAML(Yang, 2007) version 4.9h100 using input divergence time (109.8 - 124.4 MYA) between *Psidium guajava* and *Vitis vinifera* from TimeTree (Kumar et al., 2022). The overrepresentation of GO ids in the genome-specific GO term databases built using Blast2GO functional annotation was then tested using the gene lists. The visualization of the clusters was done in OrthoVenn’s(Wang et al., 2015) Cluster-Venn web module. Hypergeometric testing was done implementing extra conditioning on GO word. Hierarchical structure was used to identify overrepresentation (P 0.05) and results were plotted using R. AS 17,395 predicted genes were subjected to phylostratiography analyses using phylostratr(Arendsee et al., 2019). The focal species was set as ‘120290’ for *Psidium guajava* and the protein file was replaced by the proteome of *Psidium guajava* cv Allahabad Safeda, and default options were used. Genes found only in the *Psidium guajava* are considered orphan and assigned to the phylostratum.

### Lycopene estimation

For estimating Lycopene (mg/100 gm), 2 gm guava pulp was crushed in acetone till it became colourless. Carotenoid pigments were separated with Petroleum ether AR, 60-80°C and sodium sulphate. The optical density was noted at 503 nm and lycopene content was calculated using following formula

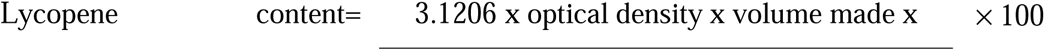

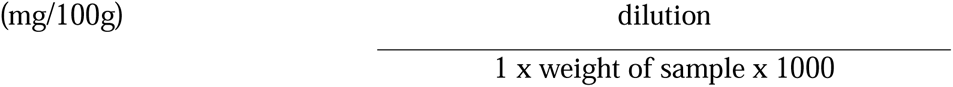

### Candidate gene screening for pulp colouration

Re-sequencing raw reads obtained of 19 variable flesh guava genotypes were quality checked by FastQC, v0.11.8 (http://www.bioinformatics.babraham.ac.uk/projects/fastqc). Trimmomatic version 0.39(Bolger et al., 2014) was used to remove low-quality reads and adapter sequences. Bowtie2(Langmead and Salzberg, 2012) was used for indexing the reference genome and mapping clean reads to indexed genome assembly. All the carotenoid pathway genes were screened for any variation (Insertion, deletion, SNP) amongst the pink and white coloured genotypes with Integrative Genomics Viewer (IGV)(Robinson et al., 2011).

### Validation for polymorphism and Protein Protein Interaction in *Phytoene Synthase* 2

Primers designed with Primer3 version 0.40 (http://primer3.sourceforge.net) spanning the InDel were tested on all guava genotypes with PCR profile - 95°C for 3 min (initial denaturation), 95°C for 30 s (denaturation), 55°C for 30 s (annealing) and 72°C for 1 min (elongation) for 35 cycles and a final elongation step at 72°C for 5 min in 10μl reaction. ∼7ng DNA was used as template with .5μM forward and reverse primers, 10 mg/ml BSA and 10mg/ml PVP in GoTaq® Green master mix (Promega Biotech India Pvt. Ltd., New Delhi, India). 3.5% agarose gel was used for resolving the amplified products and were visualised with AlphaView software in Alphaimager HP gel documentation system (ProteinSimple, San Jose, CA, United States). *In-silico* protein-protein interaction of PSY2 was done using STRING (version 10.0; http://string-db.org) followed by visualization of the top interacting proteins.

### Genome wide association for pulp colour

For genotyping by sequencing genomic DNA was extracted by CTAB method from 67 white fleshed genotypes viz. 2/2, AC 1-4, AC 1-5, AC 2-1, AC 2-3, AC 3-4, AC 5-2, AC 5-5, AC 5-6, AC 5-7, AC 6-2, AC 6-4, AC8-2, AC 8-4, AC 8-5, AC 8-6, AC 9, AC 10-5, AC 10-7, AC 10-8, AC 11-9, AC 14-4, AC 18-4, AICRP 10-5, Arka Amulya, AS, AS 1-6, AS 1-7, AS 2-8, AS 5-8, AS 7-3, AS 7-7, AS 7-8, AS 11-3, AS 12-2, AS 13-2, AS 15-3, Barf Khana, BDS, Behat Coconut, BP-1, BP-2, CISH-G1, CISH-G5 (Lalima), CISH-G6, G-Bilas, GVP, HB 2-6, HB 4-8, HB 5-8, HB 9-7, Hisar Safeda, KG, KMG, L-49, Pant Prabhat, Punjab Safeda, R2P4, Sabir, Sabir 2, Sabir 3, Seedless, Shweta, Strawberry guava, Thai guava, TS-1, VNR-Bihi and 15 Pink fleshed genotypes AICRP 10-4, Arka Kiran, HB 6-3, HB 9-2, HB 17-16, HB-88, Hisar Surkha, Lalit, Lemon guava, Pear Shaped, Portugal, Punjab Bold, Punjab Kiran, Punjab Pink and, Taiwan pink. DNA isolated from 82 guava genotypes was subjected to ddRAD sequencing where EcoRI and MseI restriction enzymes were used. Clean and trimmed ddRAD reads mapped to indexed AS genome assembly computed with Freebayes(Garrison and Marth, 2012) called SNPs that were filtered with VCFtools(Danecek et al., 2011) and TASSEL(Bradbury et al., 2007). Association between the genotypic data and phenotypic data was made with GAPIT3(Wang and Zhang, 2021).

## Supporting information

Supplementary Datafile1_22

## Acknowledgements

Authors thank Dr. John Edward Bowers, PGML, UGA, Georgia, USA for useful discussions on genome evolution. Authors also thank field men Mr. Rajinder Singh, Mr. Ranjit Singh, Mr. Jagpreet Singh and Mr. Rustum Singh for providing the support for the successful conduct of the study.

## Contributions

AM, MISG, PC, NKA, ISY – conceived the idea and obtained the funding. MISG, RSB, NKA, DS-procured and maintained the germplasm in Orchards. AM, ST, MJ, DS, RSB, NKA – Collected the leaves and fruit samples. AM, ST, MJ – Extracted the DNA from leaves. OKM, VBRL – Genome assembly with MaSuRCA. ST, AM – Gene prediction and annotation. ST – Lycopene estimation. ST, AS, AHP, GSD, ISY - conducted the bioinformatics analysis. AS, ST, GSD - graphical representations. AM, ST, AS, PC – drafted the manuscript. AM, ST, AS, AHP, GSD – manuscript preparation and revisions

## Data availability statement

Short read Illumina sequencing data, PacBio reads and BioNanoMaps have been deposited in NCBI under Bioproject PRJNA6289240. Genome re-sequencing Illumina reads from 20 genotypes are deposited under Bioproject PRJNA565883. Guava genome assembly deposited with NCBI has accession no. VYWX00000000 and is deposited under Bioproject PRJNA564981. Eighty-two genotypes ddRAD data submission to NCBI has BioProject ID PRJNA1023166.

## Supplementary Information

**Figure S1.** Summary of conserved orthologous genes in the assembled guava genome with BUSCO. The figure depicts the guava genome assembly’s completeness using the BUSCO (Benchmarking Universal Single-Copy Orthologs) study. The X-axis shows the proportion of full or fragmented BUSCO among 2121 eudicot genes.

**Figure S2.** QUality ASsessment Tool (QUAST) evaluation of genome assembly A) The GC content distribution of contigs B) The cumulative length of the contigs C) GC content distribution D) Contig length contributing to total assembly length.

**Figure S3.** LTR Assembly Index (LAI) of guava genome assembly.

**Figure S4.** GenomeScope K-mer frequency distribution plots determine k-mer based genome size and heterozygosity of the guava genome assembly.

**Figure S5.** Microsatellite Distribution & Motif Abundance: The figure displays the distribution pattern, abundance and different types of microsatellites obtained using the Krait SSR mining software. The motif abundance chart shows the relative abundance of different SSR motif in the genome.

**Figure S6.** Top 20 Gypsy LTR Families: The figure shows the top 20 Gypsy long terminal repeat (LTR) identified in the guava genome, ranked by copy number. The data points represent the whole genome percentage of the LTR and its corresponding copy number. The names of each LTR family are shown along the x-axis.

**Figure S7.** Top 20 Copia LTR Families: The figure shows the top 20 Copia long terminal repeat (LTR) identified in the guava genome, ranked by copy number. The data points represent the whole genome percentage of the LTR and its corresponding copy number. The names of each LTR family are shown along the x-axis.

**Figure S8. A.** Chromosome wise **B.** whole genome - distribution of insertion time shows that Gypsy and Copia in guava appeared from 0 to 5 million years ago **C.** Phylogenetic analysis of intact LTRs (containing GAG, PROT, RH, RT, INT domains).

**Figure S9. Ks distributions of gene pairs derived from different modes of duplication.** WGD: whole-genome duplication, DSD: dispersed duplication, PD: proximal duplication, TRD: transposed duplication.

**Figure S10. A. UpSet plot displays the unique and shared orthologous clusters among the species.** The left horizontal bar chart represents the number of orthologous clusters per species, while the right vertical bar chart indicates the number of orthologous clusters shared among the species. The intersecting lines illustrate the sets of shared clusters. B. **The heatmap shows the number of overlapping clusters between each pair of species.** A higher overlap is exhibited between the Species belonging to Myrtacea family.

**Figure S11**. **Functional Enrichment Analysis of Single Copy Clusters:** The figure showcases the results of a functional enrichment analysis performed on Single copy clusters from Orthogroup analysis. Enriched Gene Ontology (GO) terms with a corrected P value < 0.001 are displayed, with the color of circles indicating the statistical significance of the enrichment. The size of the circles corresponds to the number of genes associated with each GO term.

**Figure S12. Phylogenetic tree of 11 plant species**. Expansion and contraction of gene families is indicated by blue and pink colour, respectively in the pie chart.

**Figure S13. Functional Enrichment Analysis of Expanded and Contracted Gene Families:** The figure showcases the results of a functional enrichment analysis performed on expanded gene families (EGFs) triangle and contracted gene families (CGFs) circles. Enriched Gene Ontology (GO) terms with a corrected P value < 0.001 are displayed, with the color of circles indicating the statistical significance of the enrichment. The size of the circles and triangles correspond to the number of genes associated with each GO term.

**Figure S14**. **A phylogenetic tree based on single-copy genes illustrates the evolutionary relationship and distances among the 11 species**: The placement of *P. guajava*, *E. grandis*, *M. polymorpha* and *S. grande* within the Myrtaceae family in a monophyletic group, while *P. granatum* was placed in the Myrtales order. The bar plots represent the number of genes in cluster specific to each species.

**Figure S15. A.** Phylogenetic analysis of TPS protein sequences of guava **B.** Heat map of differentially expressed terpene synthases in Leaf, Flower, and Fruit.

**Figure S16. Dot plots of the Synteny between Psidium guajava cv Allahabad Safeda (y-axis) and genomes of Myrtacea family A.** Eucalyptus grande **B.** Syzygium grande **C.** Corymbia citridora **D.** Metrosideros polymorpha **E.** Rhodamnia argentea **F.** Psidium guajava cv. New Age. Synteny amongst all the myrtale genomes displays similar patterns.

**Figure S17. Dot plots of the Synteny between *Psidium guajava* cv Allahabad Safeda and genomes of A.** *Populus tricocarpa* **B.** *Vitis vinifera*

**Figure S18. Distribution of synonymous substitution (Ks) of syntenic orthologues**

**Figure S19. Protein-Protein Interaction Network of PSY2 Gene in *Psidium guajava* c.v Allahabad Safeda.** The figure represents the Protein-Protein Interaction (PPI) network of the PSY2 (pgpauas08401) in *Psidium guajava*. Each node in the network corresponds to a protein, and the edges between the nodes indicate their interactions. The PSY2 gene is highlighted in the network, demonstrating its connections and potential functional partners.

**Table S1.** Short read and long read sequences of Allahabad Safeda

**Table S2.** Repeats content statistics of reference genome assembly

**Table S3.** Duplications statistics of reference genome assembly

**Table S4.** Distribution of 63 classes of 875 Transcription factors (TFs) in Allahabad Safeda genome

**Table S5.** Distribution of 23 classes of 325 Transcription regulators (TRs) in Allahabad Safeda genome

**Table S6.** Classification and statistics of Terpene synthases (TPS) in guava

**Table S7.** Lycopene synthesis pathway candidate genes and isoforms

**Table S8.** *In-silico* structural variations in Phytoene synthase 2 in white vs pink pulp guava genotypes with re-sequencing data analysis

**Table S9.** Primer sequence for deletion scoring in Phytoene synthase 2 of pink pulp genotypes

**Table S10.** Genome wide single nucleotide associations for pink pulp colour with ddRAD

**Table S11.** *In silico* Phytoene Synthase 2 protein interaction analysis with guava proteins

**Supplementary Datafiles 1-22**

