## Supplementary material for "Guava *cv.* Allahabad Safeda Chromosome scale assembly and comparative genomics decodes breeders’ choice marker trait association for pink pulp colour": Figure S1_S19; Table S1_S11: Guava_Pulp_AM_Suppl.docx

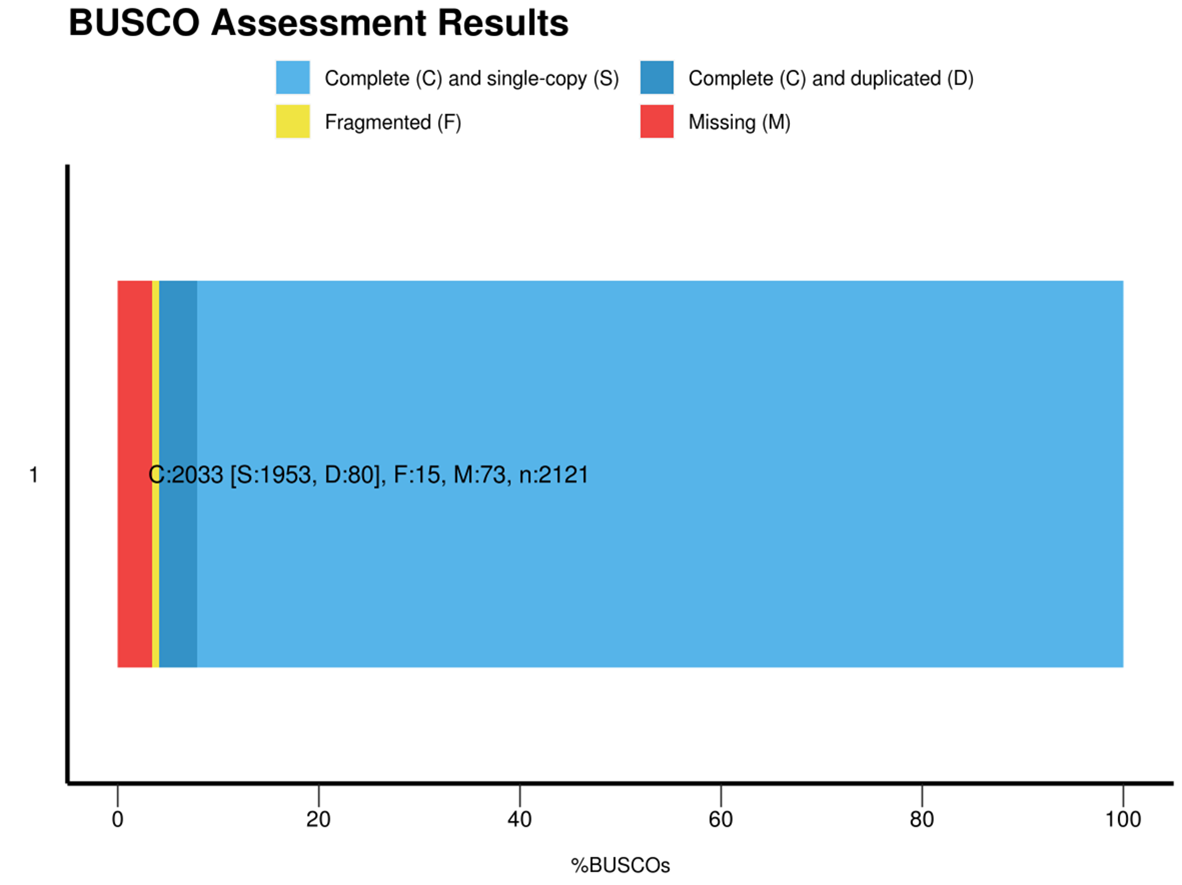

**Figure S1. Summary of conserved orthologous genes in the assembled guava genome with BUSCO.** The figure depicts the guava genome assembly's completeness using the BUSCO (Benchmarking Universal Single-Copy Orthologs) study. The X-axis shows the proportion of full or fragmented BUSCO among 2121 eudicot genes.

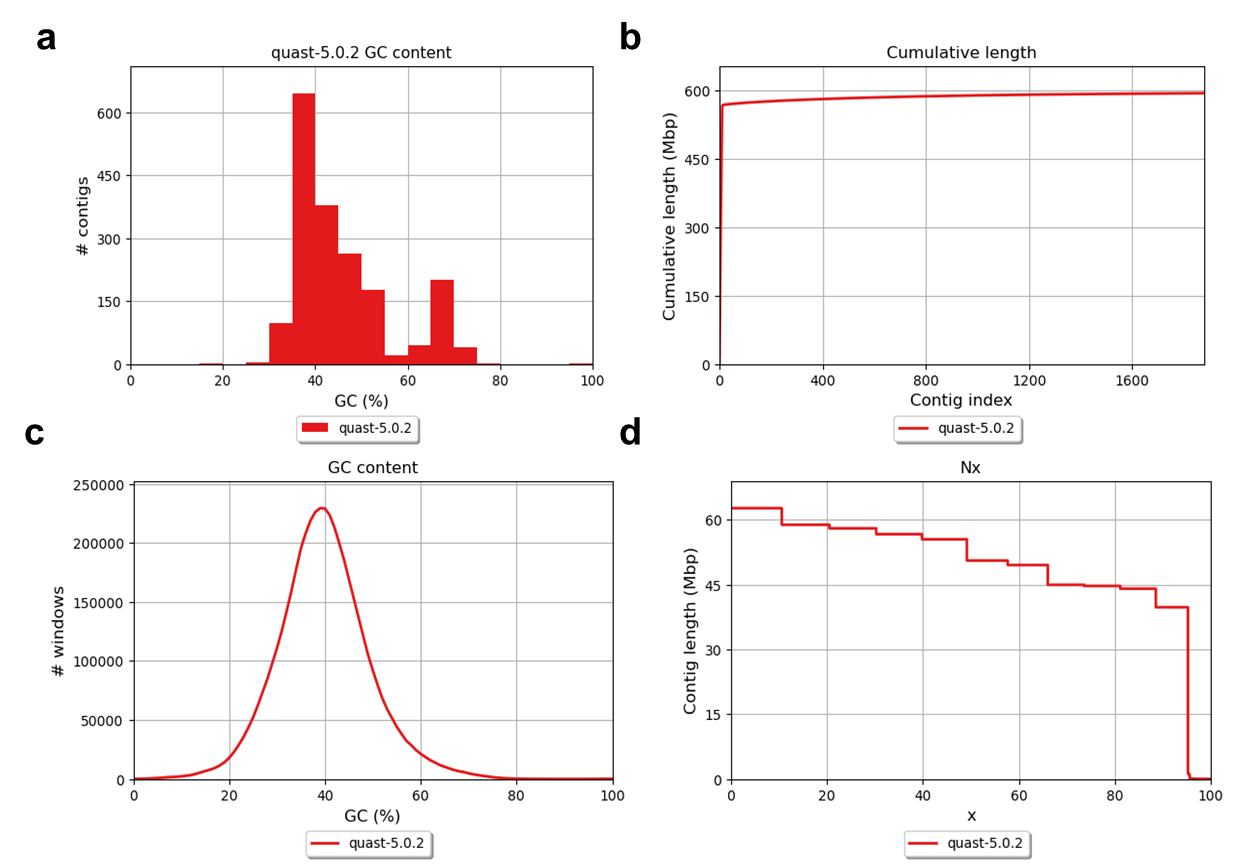

**Figure S2. QUality ASsessment Tool (QUAST) evaluation of genome assembly A)** The GC content distribution of contigs **B)** The cumulative length of the contigs **C)** GC content distribution **D)** Contig length contributing to total assembly length.

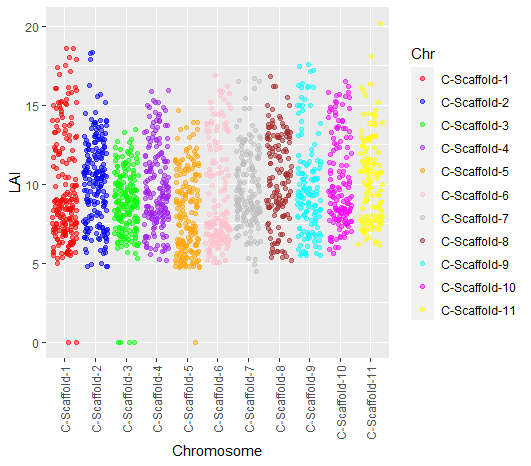

**Figure S3 LTR Assembly Index (LAI) of guava genome assembly**.

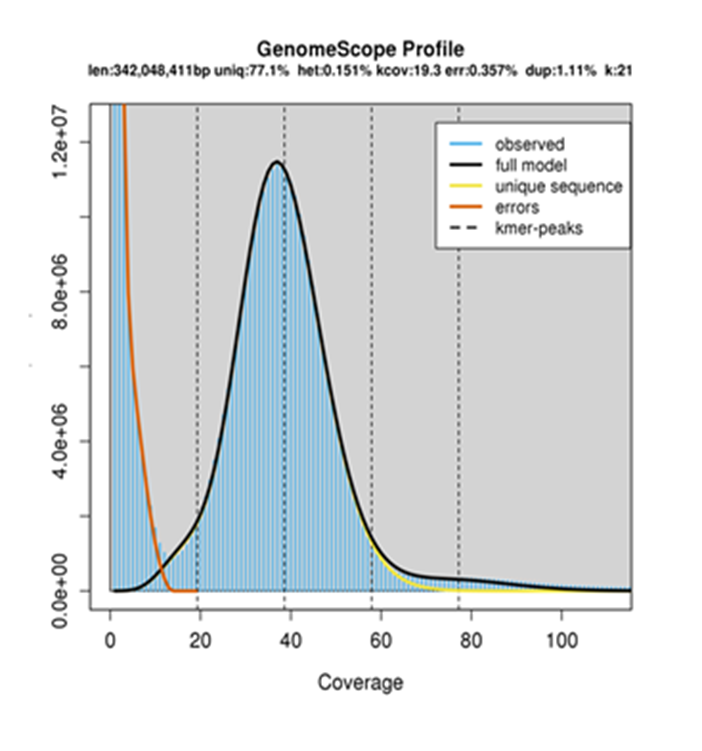

**Figure S4 GenomeScope K-mer frequency distribution plots** determine k-mer based genome size and heterozygosity of the guava genome assembly.

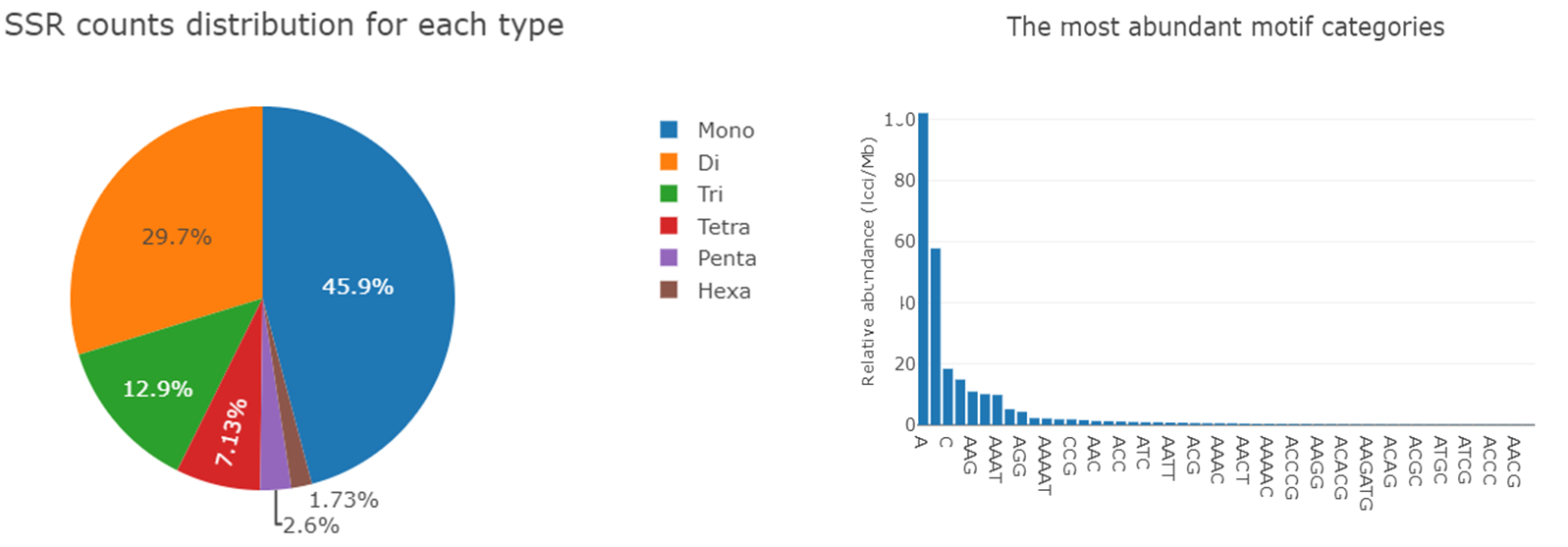

**Figure S5 Microsatellite Distribution & Motif Abundance:** The figure displays the distribution pattern, abundance and different types of microsatellites obtained using the Krait SSR mining software. The motif abundance chart shows the relative abundance of different SSR motif in the genome.

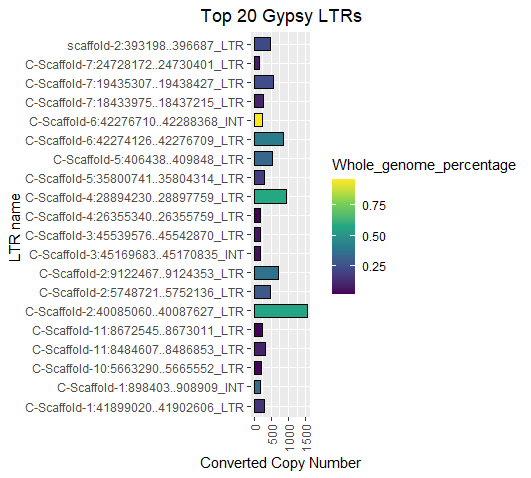

**Figure S6 Top 20 Gypsy LTR Families:** The figure shows the top 20 Gypsy long terminal repeat (LTR) identified in the guava genome, ranked by copy number. The data points represent the whole genome percentage of the LTR and its corresponding copy number. The names of each LTR family are shown along the x-axis.

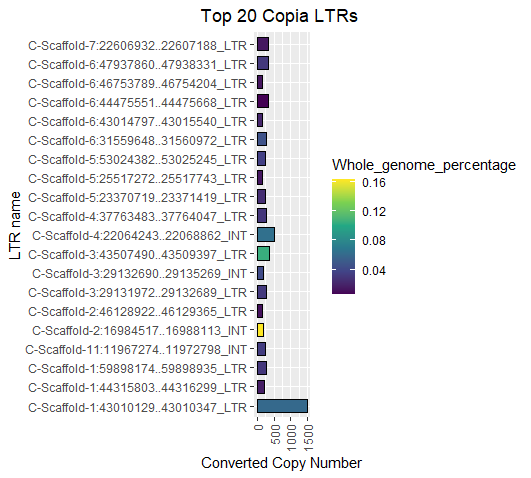

**Figure S7 Top 20 Copia LTR Families:** The figure shows the top 20 Copia long terminal repeat (LTR) identified in the guava genome, ranked by copy number. The data points represent the whole genome percentage of the LTR and its corresponding copy number. The names of each LTR family are shown along the x-axis.

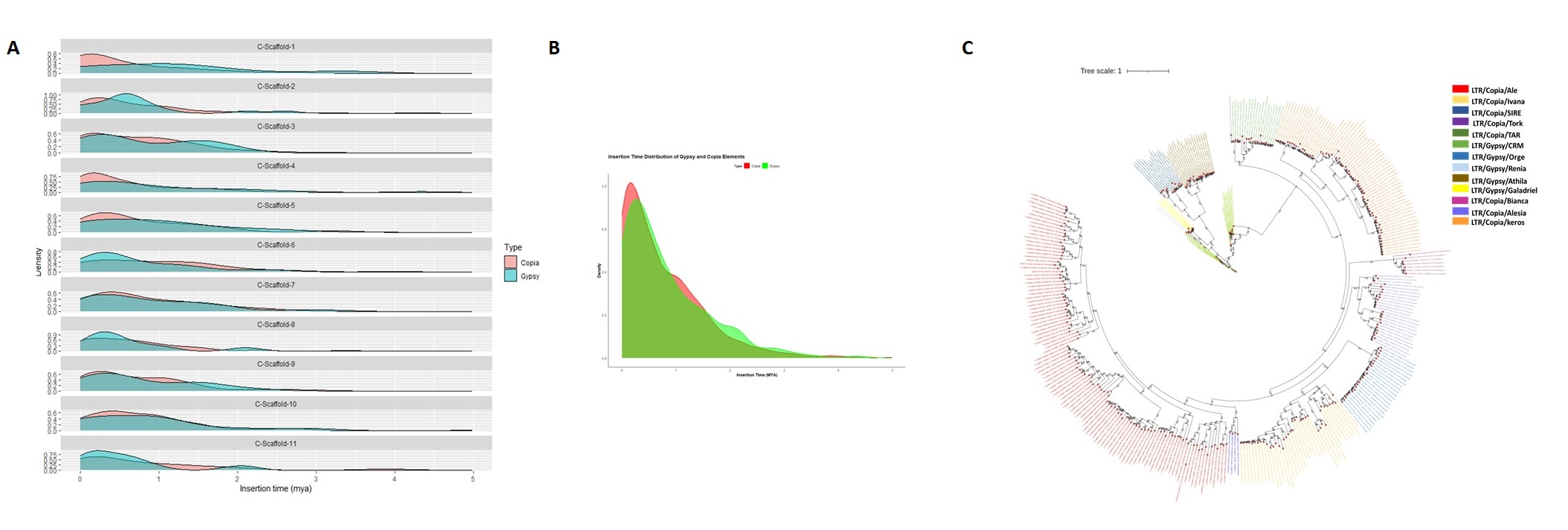

**Figure S8**. A. Chromosome wise B. whole genome - distribution of insertion time shows that Gypsy and Copia in guava appeared from 0 to 5 million years ago C. Phylogenetic analysis of intact LTRs (containing GAG, PROT, RH, RT, INT domains).

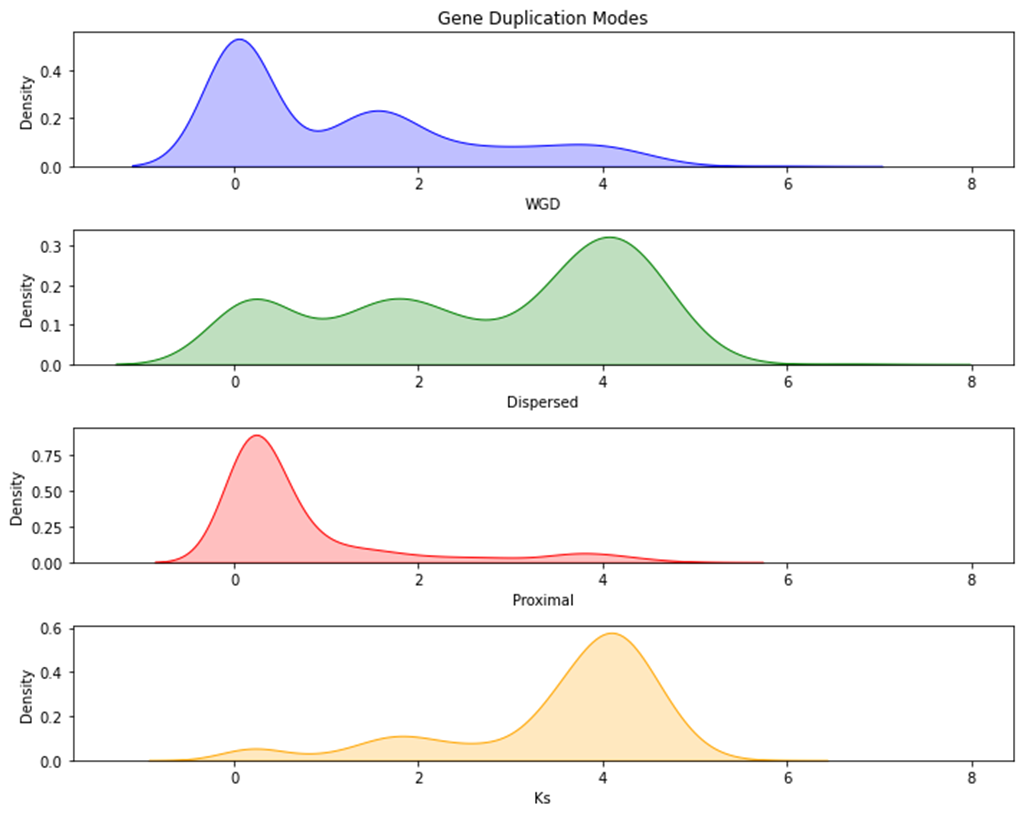

**Figure S9: Ks distributions of gene pairs derived from different modes of duplication.** WGD: whole-genome duplication, DSD: dispersed duplication, PD: proximal duplication, TRD: transposed duplication.

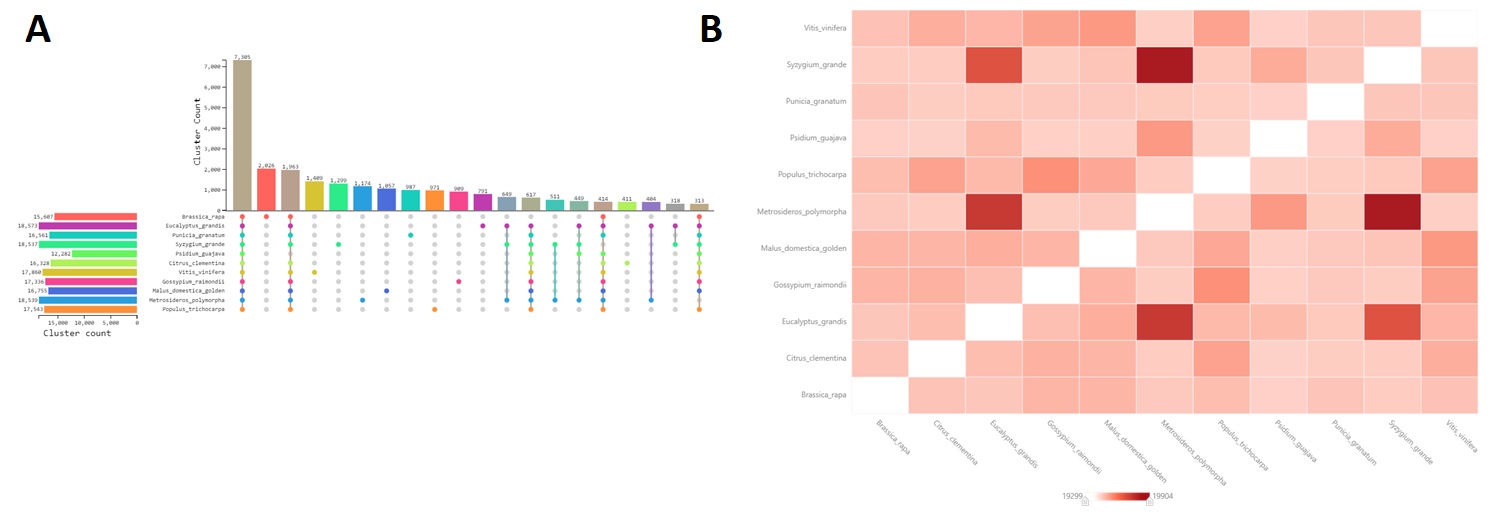

**Figure S10 A) UpSet plot displays the unique and shared orthologous clusters among the species.** The left horizontal bar chart represents the number of orthologous clusters per species, while the right vertical bar chart indicates the number of orthologous clusters shared among the species. The intersecting lines illustrate the sets of shared clusters. B) **The heatmap shows the number of overlapping clusters between each pair of species.** A higher overlap is exhibited between the Species belonging to Myrtacea family.

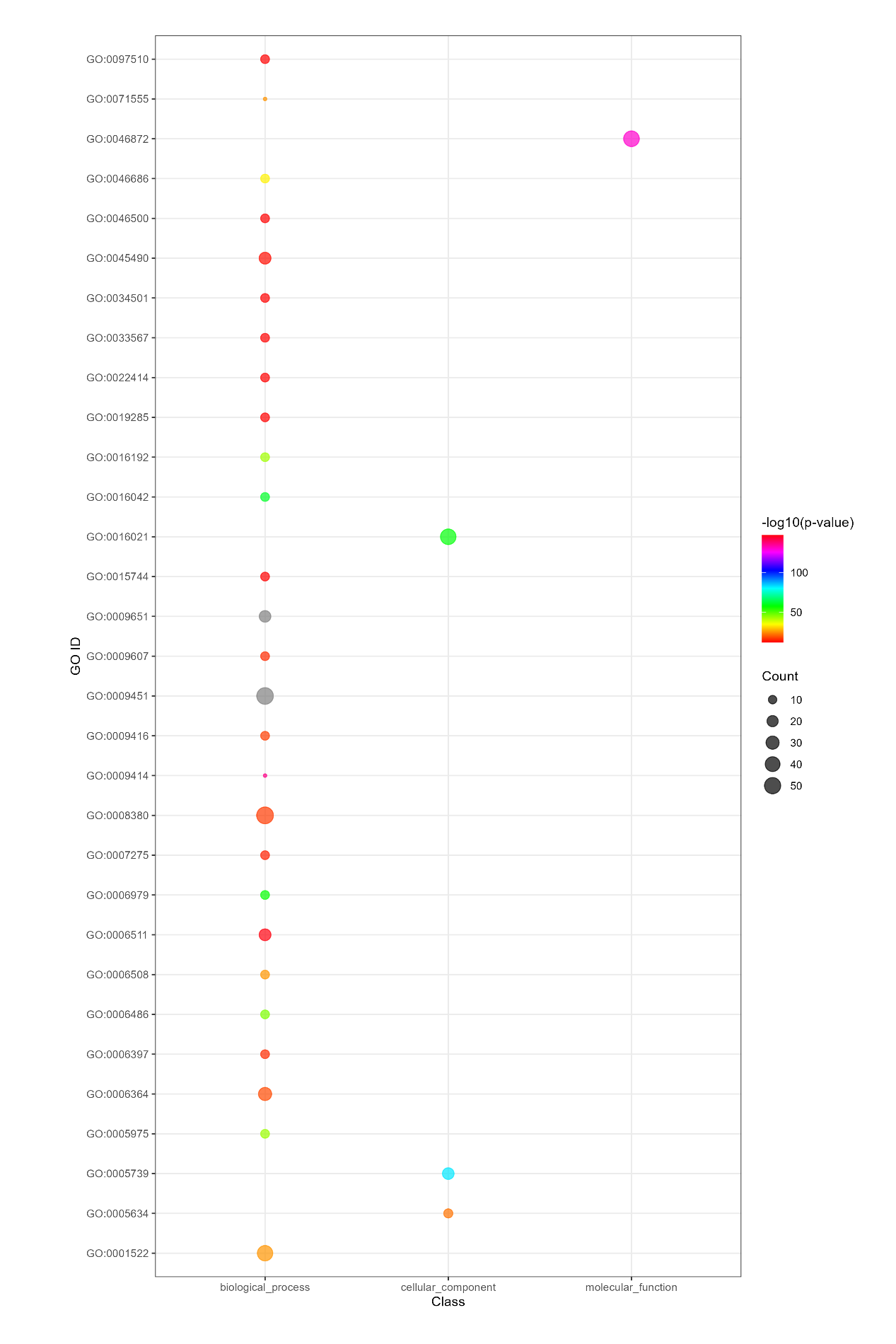

**Figure S11 Functional Enrichment Analysis of Single Copy Clusters:** The figure showcases the results of a functional enrichment analysis performed on Single copy clusters from Orthogroup analysis. Enriched Gene Ontology (GO) terms with a corrected P value < 0.001 are displayed, with the color of circles indicating the statistical significance of the enrichment. The size of the circles corresponds to the number of genes associated with each GO term.

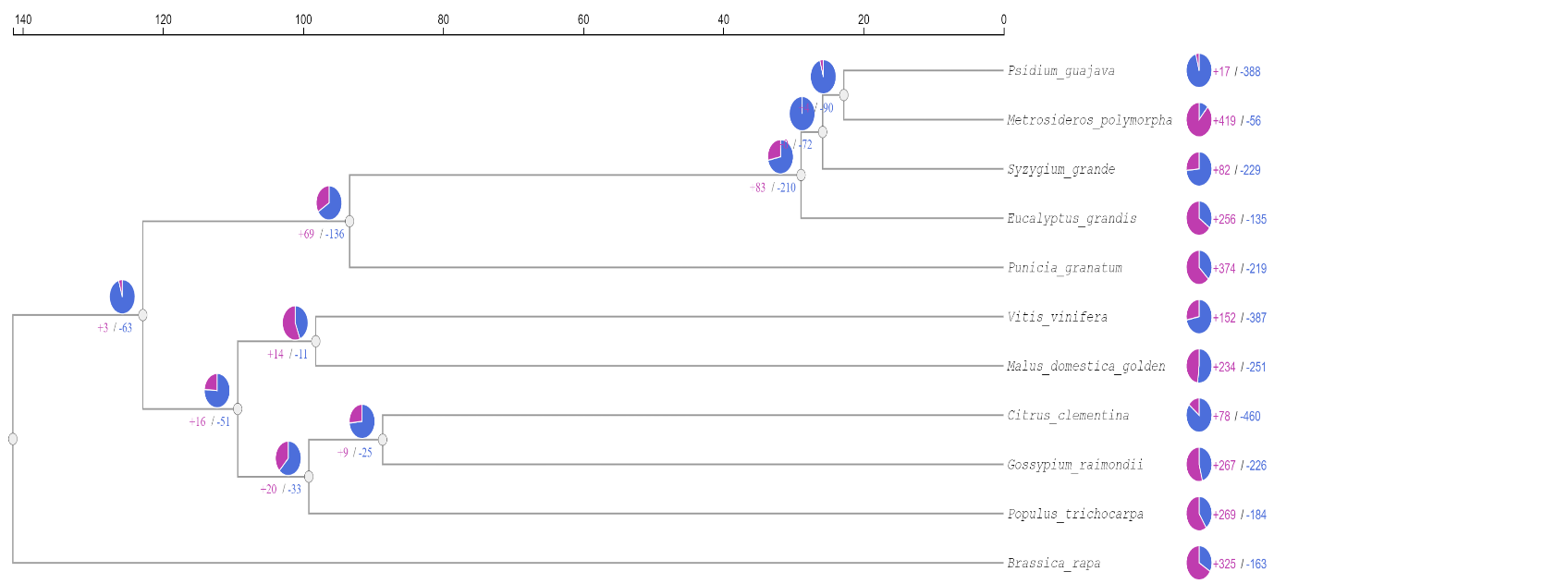

**Figure S12: Phylogenetic tree of 11 plant species**. Expansion and contraction of gene families is indicated by blue and pink colour, respectively in the pie chart.

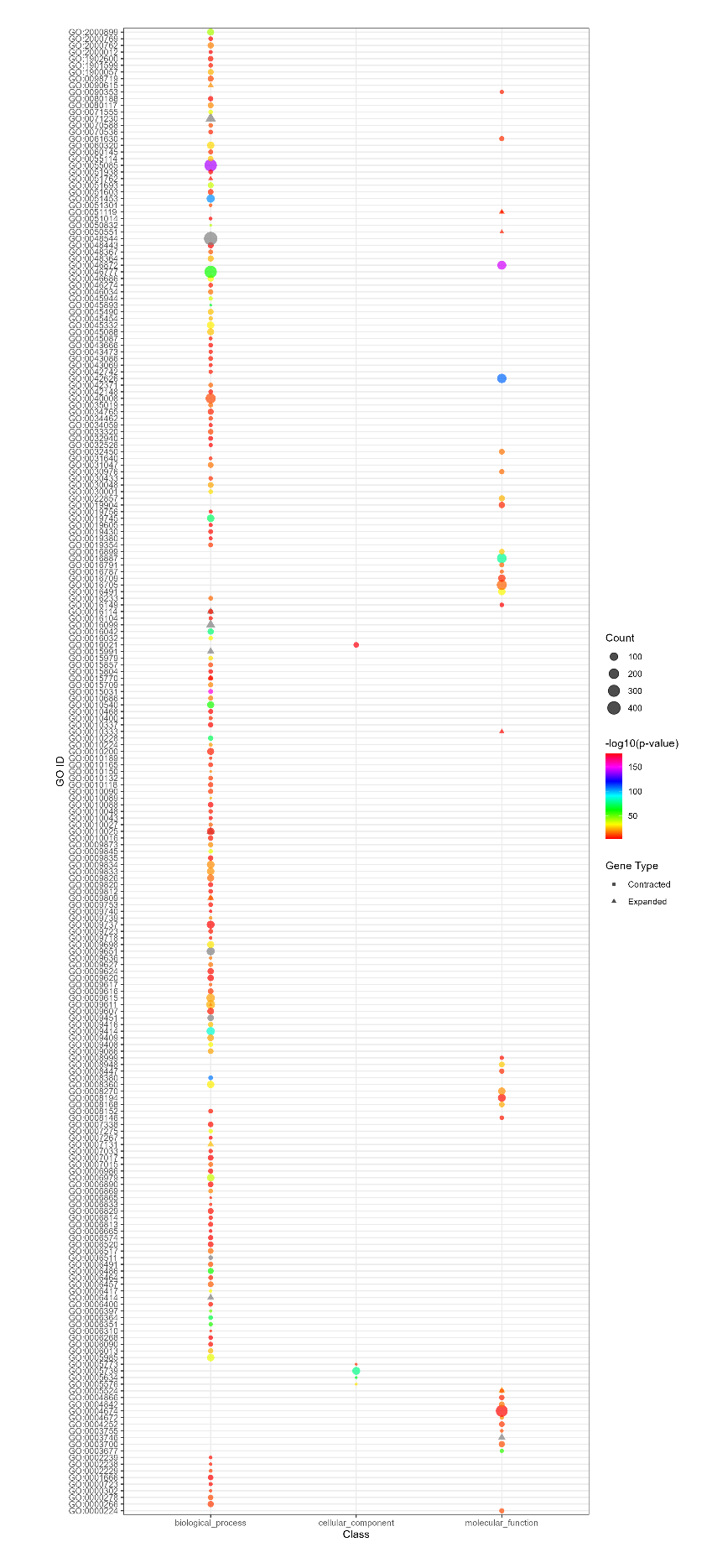

**Figure S13 Functional Enrichment Analysis of Expanded and Contracted Gene Families:** The figure showcases the results of a functional enrichment analysis performed on expanded gene families (EGFs) triangle and contracted gene families (CGFs) circles. Enriched Gene Ontology (GO) terms with a corrected P value < 0.001 are displayed, with the color of circles indicating the statistical significance of the enrichment. The size of the circles and triangles correspond to the number of genes associated with each GO term.

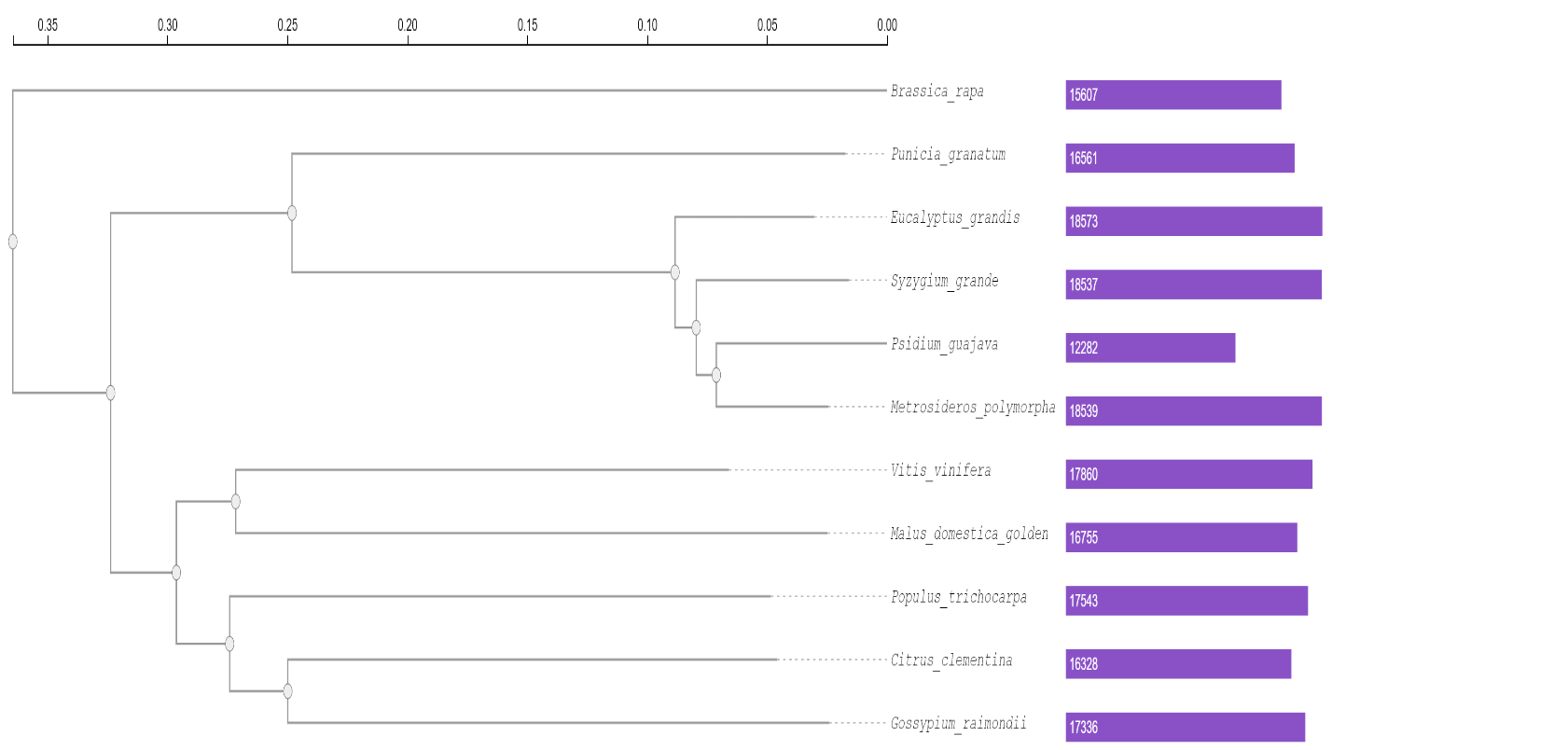

**Figure S14 A phylogenetic tree based on single-copy genes illustrates the evolutionary relationship and distances among the 11 species.** The placement of *P. guajava*, *E. grandis*, *M. polymorpha* and *S. grande* within the Myrtaceae family in a monophyletic group, while *P. granatum* was placed in the Myrtales order. The bar plots represent the number of genes in cluster specific to each species.

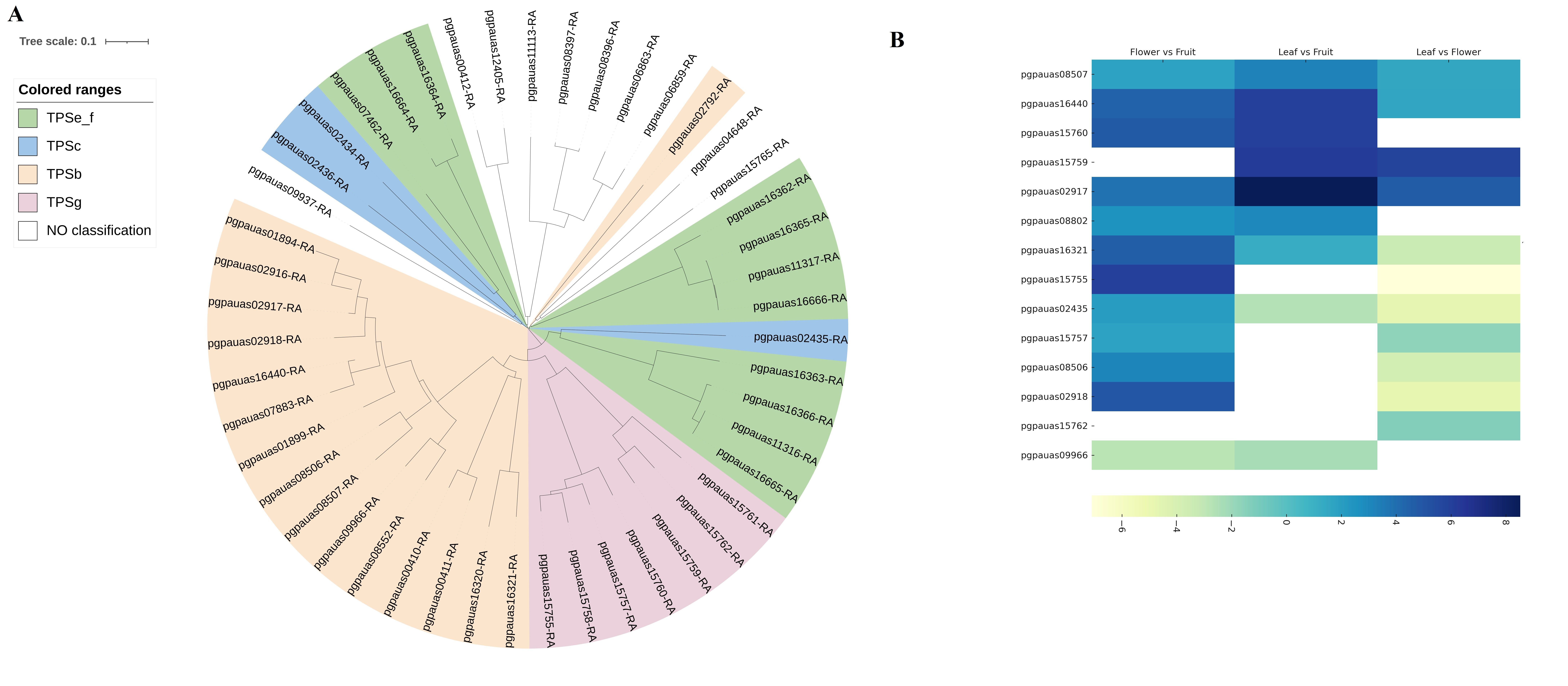
**Figure S15**: A. Phylogenetic analysis of TPS protein sequences of guava B. Heat map of differentially expressed terpene synthases in Leaf, Flower, and Fruit

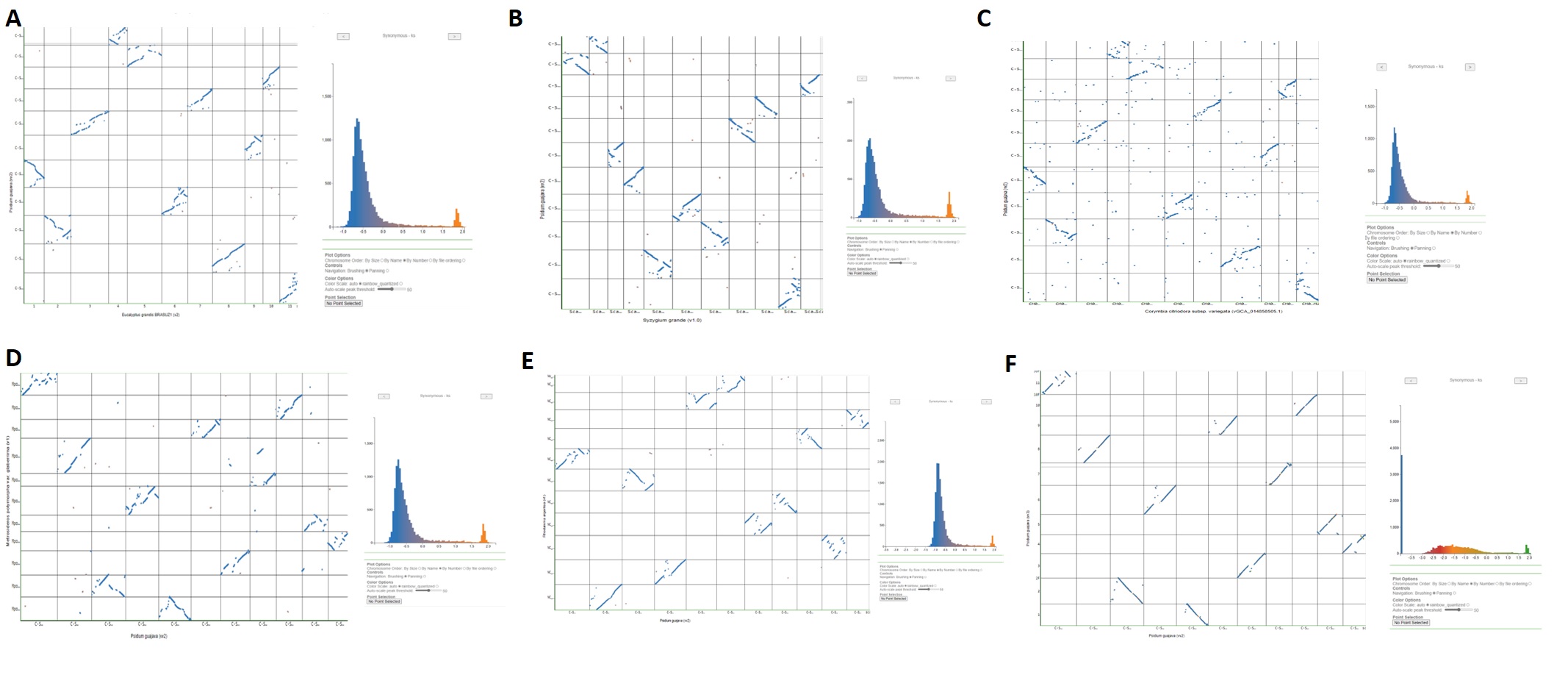

**Figure S16 Dot plots of the Synteny between *Psidium guajava* cv Allahabad Safeda (y-axis) and genomes of Myrtacea family A)** *Eucalyptus grande* **B)** *Syzygium grande* **C)** *Corymbia citridora* **D)** *Metrosideros polymorpha* **E)** *Rhodamnia argentea* **F)** *Psidium guajava* cv. New Age. Synteny amongst all the myrtale genomes displays similar patterns.

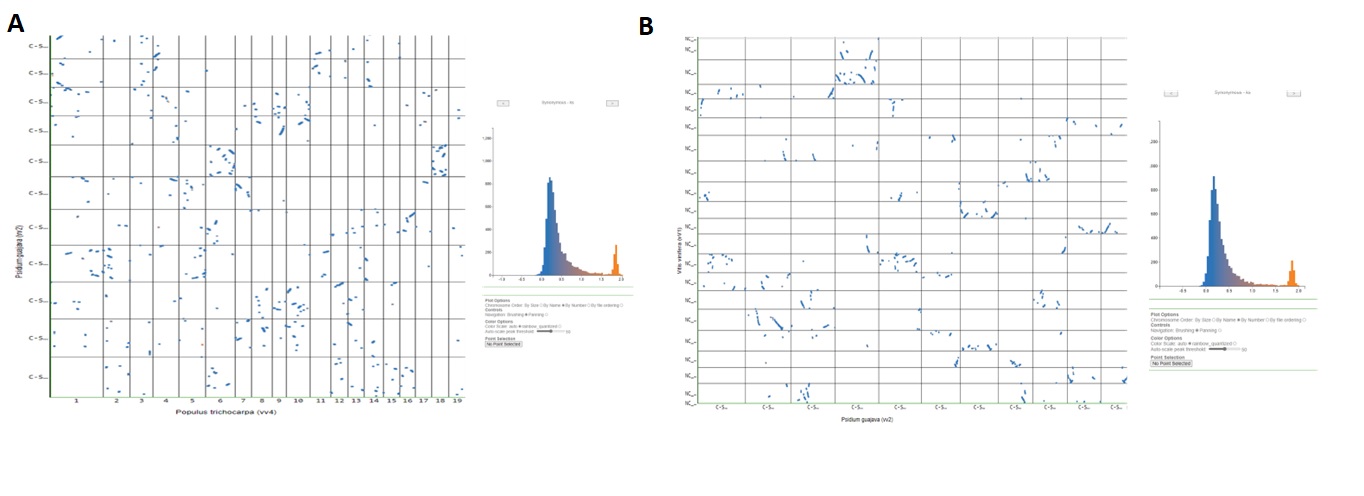

**Figure S17 Dot plots of the Synteny between *Psidium guajava* cv Allahabad Safeda and genomes of A)** *Populus tricocarpa* **B)** *Vitis vinifera*

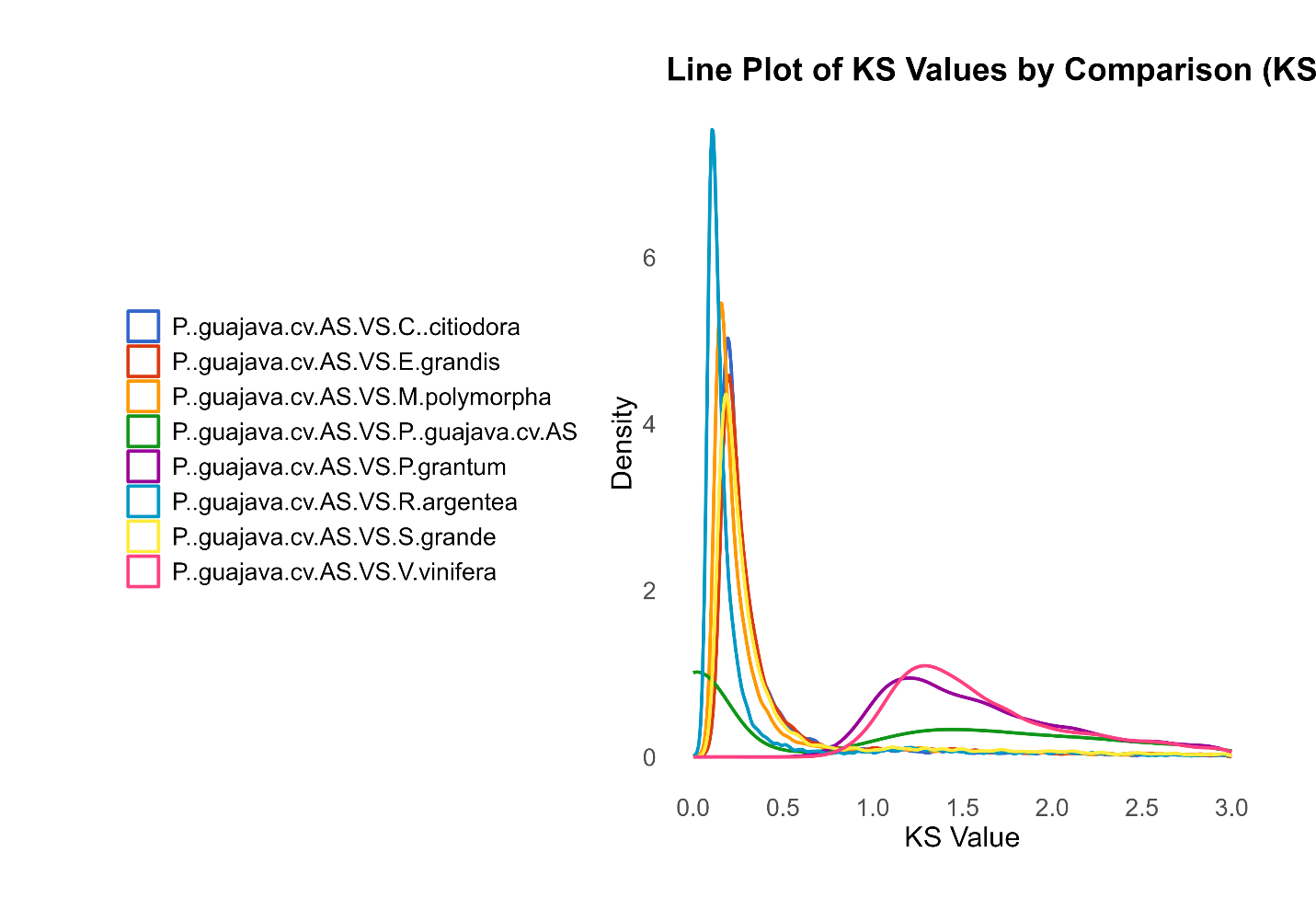

**Figure S18 : Distribution of synonymous substitution (Ks) of syntenic orthologues**

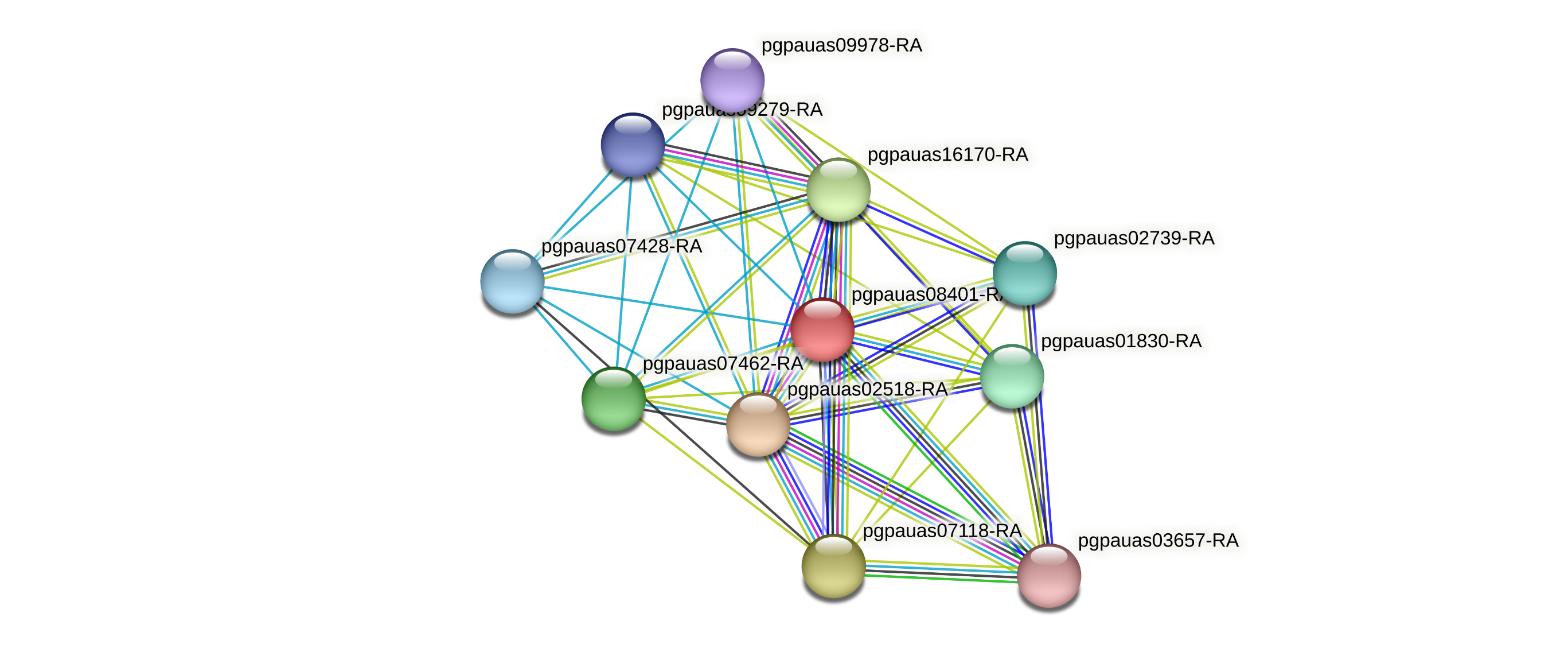

**Figure S19 Protein-Protein Interaction Network of PSY2 Gene in *Psidium guajava* c.v Allahabad Safeda.** The figure represents the Protein-Protein Interaction (PPI) network of the PSY2 (pgpauas08401) in *Psidium guajava*. Each node in the network corresponds to a protein, and the edges between the nodes indicate their interactions. The PSY2 gene is highlighted in the network, demonstrating its connections and potential functional partners.

**Table S1:** Short read and long read sequences of Allahabad Safeda

| **Illumina HiSeq x 10 – 150bp x 2 Short reads** | | | |
| --- | --- | --- | --- |
| Fragment insert size (bp) | 300 | 500 | 800 |
| PE- reads | 827,935,470 | 402,367, 298 | 471,789,730 |
| Number of bases (GB) | 124.190 | 60.355 | 70.768 |
| **PacBio SEQUEL >10 Kb Long reads** | | | |
|  | SMRT1 | SMRT2 | **Total / Average** |
| Number of bases (GB) | 10.09 | 8.57 | 18.66 |
| CCS- Reads | 788,685 | 794,336 | 1,583,021 |
| Mean Read length | 12,804 | 10,794 | 11,799 |
| Read N50 | 20,649 | 17,664 | 19,157 |
| Mean Insert Length | 11,943 | 10,137 | 11,040 |
| Insert N50 | 18,911 | 16,255 | 17,583 |

**Table S2:** Repeats content statistics of reference genome assembly

| **Class of repeat elements** | **Length occupied (bp)** | **Genome Per centage** |
| --- | --- | --- |
| LTR elements | 99518598 | 16.74 |
| Gypsy | 54127629 | 9.1 |
| Copia | 43430503 | 7.31 |
| Non-LTR elements |  |  |
| SINEs | 106988 | 0.02 |
| LINEs | 7727074 | 1.3 |
| DNA elements | 39792600 | 6.69 |
| hAT-Charlie | 587816 | 0.1 |
| MULE-MuDR | 13495136 | 2.27 |
| Satellites | 2189519 | 0.37 |
| Unclassified | 81829427 | 13.76 |
| **Total** | **234192421** | **39.39** |

**Table S3:** Duplications statistics of reference genome assembly

| **Type of Duplication** | **Number of gene pairs** |
| --- | --- |
| DSD | 10158 |
| TRD | 2175 |
| PD | 1765 |
| WGD | 991 |
| TD | 0 |

**Table S4:** Distribution of 63 classes of 875 Transcription factors (TFs) in Allahabad Safeda genome

| **S.No.** | **TF class (Domain)** | **No. of TFs** | **S.No.** | **TF class (Domain)** | **No. of TFs** |
| --- | --- | --- | --- | --- | --- |
| 1. | Alfin-like | 5 | 33. | HB-KNOX | 7 |
| 2. | AP2/ERF-RAV | 1 | 34. | HB-HD-ZIP | 22 |
| 3. | AP2/ERF-ERF | 23 | 35. | HRT | 1 |
| 4. | AP2/ERF-AP2 | 10 | 36. | HSF | 1 |
| 5. | B3 | 33 | 37. | LFY | 1 |
| 6. | B3-ARF | 9 | 38. | LIM | 7 |
| 7. | BBR-BPC | 3 | 39. | LOB | 26 |
| 8. | BES1 | 6 | 40. | MADS-M-type | 2 |
| 9. | bHLH | 74 | 41. | MYB | 88 |
| 10. | bZIP | 44 | 42. | MYB-related | 64 |
| 11. | C2C2-YABBY | 4 | 43. | NAC | 31 |
| 12. | C2C2-LSD | 3 | 44. | NF-X1 | 1 |
| 13. | C2C2-CO-like | 11 | 45. | NF-YA | 7 |
| 14. | C2C2-GATA | 23 | 46. | NF-YC | 6 |
| 15. | C2C2-Dof | 12 | 47. | NF-YB | 2 |
| 16. | C2H2 | 49 | 48. | NOZZLE | 1 |
| 17. | C3H | 39 | 49. | PLATZ | 19 |
| 18. | CAMTA | 4 | 50. | RWP-RK | 6 |
| 19. | CPP | 6 | 51. | S1Fa-like | 2 |
| 20. | DBB | 4 | 52. | SBP | 15 |
| 21. | DBP | 2 | 53. | SRS | 9 |
| 22. | DDT | 4 | 54. | TCP | 6 |
| 23. | E2F-DP | 5 | 55. | Tify | 17 |
| 24. | EIL | 2 | 56. | Trihelix | 18 |
| 25. | FAR1 | 13 | 57. | TUB | 5 |
| 26. | GARP-G2-like | 29 | 58. | ULT | 1 |
| 27. | GARP-ARR-B | 6 | 59. | VOZ | 4 |
| 28. | GRARS | 11 | 60. | Whirly | 1 |
| 29. | GRF | 4 | 61. | WRKY | 51 |
| 30. | HB-PHD | 1 | 62. | zf-HD | 1 |
| 31. | HB-WOX | 4 | 63. | HB-BELL | 7 |
| 32. | HB-other | 6 |  |  | |

**Table S5:** Distribution of 23 classes of 325 Transcription regulators (TRs) in Allahabad Safeda genome

| **S.No.** | **TR class (Domain)** | **No. of TRs** |
| --- | --- | --- |
| 1. | ARID | 9 |
| 2. | AUX/IAA | 25 |
| 3. | Coactivator p15 | 3 |
| 4. | GNAT | 21 |
| 5. | HMG | 6 |
| 6. | IWS1 | 9 |
| 7. | Jumonji | 12 |
| 8. | LUG | 5 |
| 9. | MED6 | 1 |
| 10. | MED7 | 1 |
| 11. | RB | 1 |
| 12. | Rcd1-like | 5 |
| 13. | SET | 27 |
| 14. | SNF2 | 52 |
| 15. | SOH1 | 1 |
| 16. | SWI/SNF-BAF60F | 12 |
| 17. | SWI/SNF-SW13 | 3 |
| 18. | TAZ | 7 |
| 19. | TRAF | 19 |
| 20. | mTERF | 19 |
| 21. | PHD | 30 |
| 22. | Psuedo ARR-B | 2 |
| 23. | OTHERS | 55 |

**Table S6:** Classification and statistics of Terpene synthases (TPS) in guava

| **S. No.** | **TPS gene** | **Terpene type** | **Number in *P. guajava*** |
| --- | --- | --- | --- |
| 1. | TPSa | Sesquiterpene | 1 |
| 2. | TPSb | Monoterpene | 16 |
| 3. | TPSc | Diterpene | 3 |
| 4. | TPSe_f | Mono, Di, Sesqui | 11 |
| 5. | TPSg | Mono, Di, Sesqui | 7 |
| 6. | TPSh | Di | 0 |

**Table S7:** Lycopene synthesis pathway candidate genes and isoforms

| **S.No.** | **Gene Name** | **Number of isoforms in guava genome** |
| --- | --- | --- |
| 1. | 15-cis-phytoene desaturase | 2 |
| 2. | Phytoene synthase 2 | 5 |
| 3. | 15-cis-**ζ**-carotene | 1 |
| 4. | **ζ**-carotene desaturase | 2 |
| 5. | Carotenoid cleanage dioxygenase like | 3 |
| 6. | Beta-carotene-3 hydroxylase-2 | 1 |
| 7. | Beta-carotene isomerase | 2 |
| 8. | Lycopene beta cyclase | 2 |
| 9. | Beta carotene 3 hydroxylase | 1 |
| 10. | Carotene epsilon monoxygenase | 1 |
| 11. | Geranyl geranyl pyrophosphate synthase | 1 |
| 12. | Geranyl geranyl transferase type-2 subunit alpha | 2 |
| 13. | Heterodimeric geranyl geranyl pyrophosphate synthase | 1 |
| 14. | Geranyl geranyl diphosphate reductase | 1 |
| 15. | Geranyl geranyl transferase type-1subunit beta isoform X2 | 1 |
| 16. | Geranyl geranyl transferase type-2subunit beta 1-like isoform X1 | 1 |
| 17. | Isopentenyl-diphosphate delta isomerase-I | 2 |
| 18. | Prolycopene isomerase | 5 |
| 19. | Lycopene epsilon cyclase | 2 |
| 20. | Reticulin 7-O methyltransfersae | 1 |
| 21. | CTP synthase like | 3 |
| 22. | Secoisolariciresinol dehydrogenase | 6 |
| 23. | Lutein deficient | 1 |

**Table S8:** *In-silico* structural variations in Phytoene synthase 2 in white vs pink pulp guava genotypes with re-sequencing data analysis

| **S.No.** | **Chromosome** | **Gene ID** | **Position** | **Gene feature** | **White genotypes** | **Pink genotypes** |
| --- | --- | --- | --- | --- | --- | --- |
| 1. | C-Scaffold-5 | pgpauas08401 | 26201186 | intergenic | AGAGAGAGAG | Deletion of 10 -12bp /AGAGAGAGAG |
| 2. | C-Scaffold-5 | pgpauas08401 | 26201661 | exon | T | G/T |
| 3. | C-Scaffold-5 | pgpauas08401 | 26201934 | intron | A | A/G |
| 4. | C-Scaffold-5 | pgpauas08401 | 26201955 | intron | T | C/T |
| 5. | C-Scaffold-5 | pgpauas08401 | 26201972 | intron | G | G/T |
| 6. | C-Scaffold-5 | pgpauas08401 | 26201980 | intron | A | A/G |
| 7. | C-Scaffold-5 | pgpauas08401 | 26202164 | exon | A | A/G |
| 8. | C-Scaffold-5 | pgpauas08401 | 26202280 | intron | G | G/T |
| 9. | C-Scaffold-5 | pgpauas08401 | 26202373 | intron | A | A/G |
| 10. | C-Scaffold-5 | pgpauas08401 | 26202405 | exon | G | A/G |

**Table S9:** Primer sequence for deletion scoring in Phytoene synthase 2 of pink pulp genotypes

|  |  | **PRIMER SEQUENCE** | **LENGTH** | **ANNEALING TEMPERATURE** | **PRODUCT SIZE** |
| --- | --- | --- | --- | --- | --- |
| **PSY2-InDel-1** | Forward | CCCAACTGACGGCCGATAATC | 21 | 55 ֯C | 102 |
|  | Reverse | CACATGTGAGGGACCAATCAGT | 22 |  |  |

**Table S10:** Genome wide single nucleotide associations for pink pulp colour with ddRAD

| **S.No.** | **Chromosome** | **Gene ID** | **Position** | **Gene feature** | **Gene name** |
| --- | --- | --- | --- | --- | --- |
| 1. | C-Scaffold-5 | pgpauas08401 | 26202373 | intron | Phytoene synthase 2 |
| 2. | C-Scaffold-5 | pgpauas08401 | 26202405 | exon | Phytoene synthase 2 |
| 3. | C-Scaffold-5 | pgpauas08392 | 26081386 | exon | Reticulon like protein B8 isoform X1 |

| **Gene** | **Annotation** | **node_degree** |
| --- | --- | --- |
| pgpauas01830-RA | 15-cis-phytoene desaturase, chloroplastic/chromoplastic | 8 |
| pgpauas02518-RA | geranylgeranyl pyrophosphate synthase, chloroplastic-like | 10 |
| pgpauas02739-RA | 15-cis-phytoene desaturase, chloroplastic/chromoplastic | 8 |
| pgpauas03657-RA | geranylgeranyl diphosphate reductase, chloroplastic | 5 |
| pgpauas07118-RA | solanesyl diphosphate synthase 1, chloroplastic-like | 8 |
| pgpauas07428-RA | dehydrodolichyl diphosphate synthase complex subunit nus1 | 7 |
| pgpauas07462-RA | ent-kaur-16-ene synthase, chloroplastic isoform X1 | 9 |
| pgpauas08401-RA | phytoene synthase 2, chloroplastic-like | 10 |
| pgpauas09279-RA | protein farnesyltransferase subunit beta-like | 7 |
| pgpauas09978-RA | protein farnesyltransferase subunit beta-like | 7 |
| pgpauas16170-RA | farnesyl pyrophosphate synthase 1 isoform X1 | 9 |

**Table S11:** *In silico* Phytoene Synthase 2 protein interaction analysis with guava proteins
